# Possible co-option of a VEGF-driven tubulogenesis program for biomineralization in echinoderms

**DOI:** 10.1101/554683

**Authors:** Miri Morgulis, Tsvia Gildor, Modi Roopin, Noa Sher, Assaf Malik, Maya Lalzar, Monica Dines, Lama Khalaily, Smadar Ben-Tabou de-Leon

**Affiliations:** Department of Marine Biology, Leon H. Charney School of Marine Sciences, University of Haifa, Haifa 31905, Israel.; Bionformatics Core Unit, University of Haifa, Haifa 31905, Israel.; Sagol Department of Neurobiology, University of Haifa, Haifa 31905, Israel.

**Author notes:** These authors contributed equally to this work (Miri Morgulis, Tsvia Gildor, Modi Roopin).

**Keywords:** Gene regulatory networks, biomineralization, tubulogenesis, VEGF signaling, echinoderms, evolution, morphogenesis

## Abstract

Biomineralization is the process in which living organisms use minerals to form hard structures that protect and support them. Biomineralization is believed to have evolved rapidly and independently in different phyla utilizing existing components used for other purposes. The mechanistic understanding of the regulatory networks that drive biomineralization and their evolution is far from clear. The sea urchin skeletogenesis is an excellent model system for studying both gene regulation and mineral uptake and deposition. The sea urchin calcite spicules are formed within a tubular cavity generated by the skeletogenic cells under the control the vascular endothelial growth factor (VEGF) signaling. The VEGF pathway controls tubulogenesis and vascularization across metazoans while its regulation of biomineralization was only observed in echinoderms. Despite the critical role of VEGF signaling in sea urchin spiculogenesis, the downstream program it activates was largely unknown. Here we study the cellular and molecular machinery activated by the VEGF pathway during sea urchin spiculogenesis and reveal multiple parallels to the regulation of tubulogenesis during vertebrate vascularization. Human VEGF rescues sea urchin VEGF knock-down; VEGF-dependent vesicle deposition plays a significant role in both systems and sea urchin VEGF signaling activates hundreds of genes including biomineralization and vascularization genes. Five upstream transcription factors and three signaling genes active in spiculogenesis are homologous to vertebrate factors that regulate vascularization. Overall, our findings suggest that sea urchin spiculogenesis and vertebrate vascularization diverged from a common ancestral tubulogenesis program, broadly adapted for vascularization and specifically co-opted for biomineralization in the echinoderm phylum.

**Significance statement:** The sea urchin calcite spicules and vertebrate blood vessels are quite distinct in their function, yet both have a tubular structure and are controlled by the vascular endothelial growth factor (VEGF) pathway. Here we study the downstream program by which VEGF pathway drives sea urchin spiculogenesis and find remarkable similarities to the control of vertebrate vascularization. The similarities are observed both in the upstream gene regulatory network, in the downstream effector genes and the cellular processes that VEGF signaling controls at the site of the calcite spicule formation. We speculate that sea urchin spiculogenesis and vertebrate vascularization diverged from a common ancestral tubulogenesis program that was co-opted for biomineralization in the echinoderm phylum.

## Introduction

Biomineralization is the process in which soft organic tissues use minerals to produce shells, skeletons and teeth for various functions such as protection and physical support (1). This process occurs within diverse organisms from the five kingdoms of life, bacteria, protista, fungi, animals and plants (1). Biomineralization is thought to have evolved independently and rapidly in different phyla, through the use of preexisting components and the evolution of specialized biomineralization proteins that utilize different minerals and shape them in different forms (2, 3). To understand the biological control and evolution of biomineralization it is essential to study the gene regulatory networks (GRNs) that control this process and unravel their origin. However, the structure and the function of the GRNs that control biomineralization processes were studied only for a few examples, mostly for the vertebrates’ bone and teeth formation (4, 5).

The model of GRN that controls skeletogenic cell specification in the sea urchin embryo is one of the most elaborate of its kind (6). Studies of the sea urchin larval skeleton have significantly contributed to the field of biomineralization by illuminating the pathway of mineral uptake and deposition in live embryos (7–9). Yet, the skeletogenic GRN was mostly studied at the early stages of skeletogenesis before the spicules are formed and therefore it is still unclear how the skeletogenic GRN controls spicule formation and biomineralization.

The sea urchin skeleton is made of two rods of calcite generated by the skeletogenic mesodermal (SM) cells (Fig. 1A). The SM cells ingress into the blastocoel, fuse through their filopodia and form a ring with two lateral cell clusters (red cells in Fig. 3A). In these clusters, the cells construct a syncytial cytoplasmic cord into which they secrete vesicles of calcium carbonate that form the calcite spicules (Fig. 1B-F, (7–11)). The spicule cord has a tubular structure (8, 9) and the forming calcite rods have a circular shape (Fig. 1D-F, (9)). SM cell migration, lateral cluster formation and sea urchin spiculogenesis are regulated by the vascular endothelial growth factor (VEGF) pathway (Fig. S2C, (6, 12, 13)). The sea urchin *VEGF3* gene is expressed in two lateral ectodermal domains positioned in close proximity to the SM lateral cell clusters and its receptor, VEGFR-10-Ig, is expressed exclusively in the SM cells (Fig. 3A, (12, 13). For simplicity, throughout the paper, we use the annotation *VEGF* for *VEGF3* and *VEGFR* for *VEGFR-10-Ig* (12)). VEGFR is a vital part of the echinoderm biomineralization program as it is expressed in the skeletogenic cells of all the echinoderm classes that produce larval skeletons (sea urchins and brittle stars (12, 14)) and in the skeletogenic cells of all studied adults (sea urchin and sea stars (12–15)).

**Figure 1.**
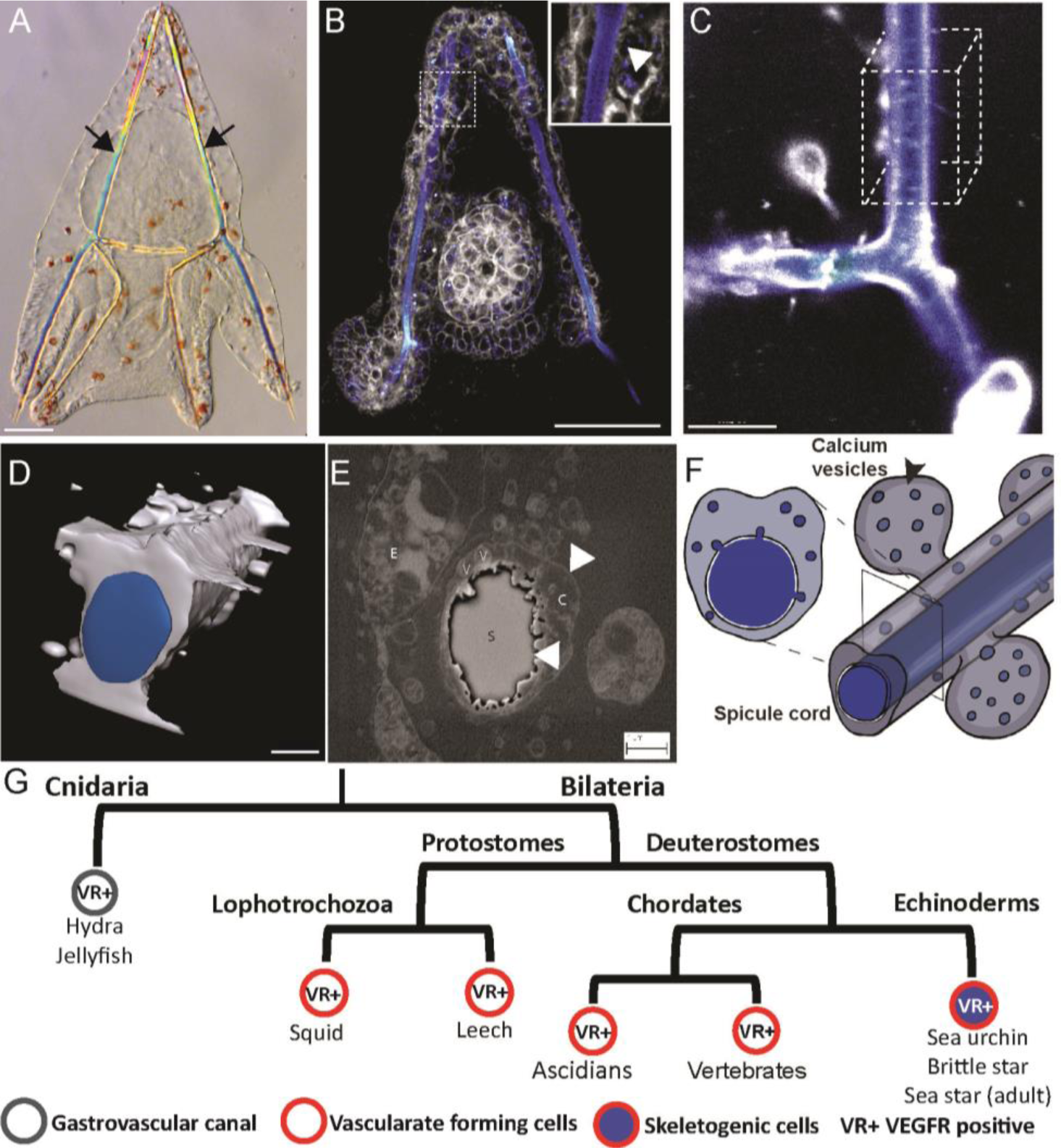
Spiculogenesis in the sea urchin embryo and VEGFR expression in tubular organs in metazoan. A. Sea urchin larva at 3dpf, showing its two calcite spicules (arrows). B, Live sea urchin embryo at 2dpf stained with the membrane tracker FM4-64 (gray) and green-calcein that binds to calcium ions (false color blue). Enlargement shows the calcium vesicles in the skeletogenic cells (arrowhead). C, A confocal image of the spicule in live embryo at 3dpf stained with blue calcein (blue) and FM4-64 (gray). D, 3D model of the spicule structure based on 50 confocal z-stacks of the cube in C. E, scanning electron micrograph of a cross section of the spicule at 3dpf showing the double membrane cytoplasmic-cord that surrounds the spicule and the vesicles inside the cord. S – spicule, C – cytoplasm, V – vesicles, E – ectodermal cell, arrows point to the membranes. Scale bars show 50 µm in A and B, scale bar shows 10µm in C, 3µm in D, 1µm in E. F, Schematic model of the spicule and vesicle secretion, showing the calcium vesicles and spicule in blue and cytoplasm in gray. G, Partial phylogenetic tree presenting VEGFR expression in cells that generate tubular structures in different phyla throughout the animal kingdom.

**Figure 3.**
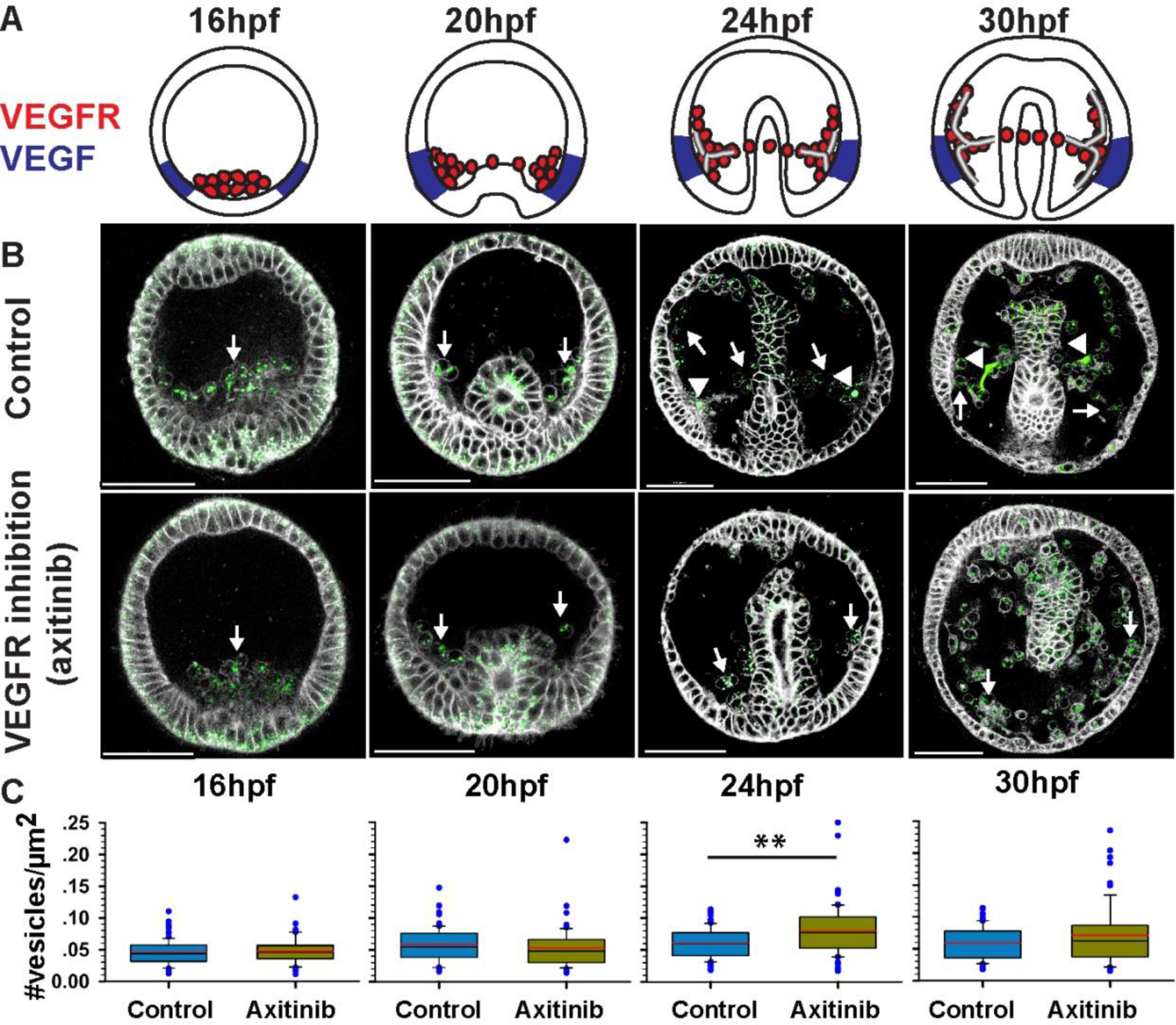
VEGF activity is essential for calcium vesicle secretion. A, schematic diagrams showing *Pl-VEGFR* expression in the skeletogenic cells (red) and *Pl-VEGF* ectodermal expression (blue) at different developmental times (similar times in A-C). B, confocal images of calcein staining (green) and FM4-64 membrane marker (white) show the presence of calcium vesicles in the SM cells (white arrows) in normal and VEGFR inhibited embryos (axitinib). Arrowheads indicate the spicules in control embryos. Scale bars are 50µm. C, vesicle number per µm^2^ in the SM cells in control and VEGFR inhibition. Each box plot shows the median (black line), average (red line) the first and the third quartiles (edges of boxes) and outliers (n=3, exact number of cells in each condition is provided in table S1).

To the best of our knowledge, VEGF signaling is not known to participate in biomineralization outside the echinoderm phylum; but it controls tubular structure formation across the animal kingdom (Fig. 1G). In vertebrates, VEGF guides the migration of hemangioblasts, the progenitors of endothelial and hematopoietic cells, and drives the formation of blood vessels during embryogenesis (vascularization), in adult ischemic tissues and in cancer (angiogenesis) (16, 17). In ascidians, VEGF signaling promotes the regeneration of blood vessels (18). Within protostomes, VEGFR is essential for blood vessel formation in the leech and the squid (19, 20). In Cnidarians VEGFR is expressed in two tubular organs: the gastrovascular canal and the tentacles (21, 22). The participation of VEGF signaling in generating tubular organs in different phyla raises the possibility that biomineralization in echinoderms is evolutionary related to these other tubulogenesis programs. Previous studies have shown that VEGFR inhibition completely abolishes spicule formation (Fig. S2C, (12, 13)), but how VEGF signaling and the skeletogenic GRN regulate biomineralization is still unclear. Here we studied the mechanisms that VEGF signaling activates during sea urchin spiculogenesis including vesicle accumulation and secretion, transcriptional targets and their function and VEGF regulation of the upstream transcription factors. We reveal intriguing similarities to the control systems that regulate vertebrate vascularization that could support a common origin of these two distinct tubulogenesis programs.

### VEGF-VEGFR recognition is conserved between human and sea urchin

600 million years of divergent evolution between vertebrates and echinoderms have generated significant differences in VEGF (Fig. S1A) and VEGFR sequences, distinctly, the sea urchin VEGFR has ten Ig domains while vertebrate VEGF receptors have seven (12). Our models of the structure of sea urchin VEGF and VEGF-VEGFR complexes show similarities to the structure of the human proteins (Fig. S1B). Yet, the question remains: are these proteins functionally similar? To test this we experimentally studied the recognition between human VEGF and sea urchin VEGFR by overexpressing human VEGF in sea urchin embryos. We injected the mRNA of one of the most abundant forms of human VEGF, *Hs-VEGFa(165*) (23) into the eggs of the Mediterranean sea urchin, *Paracentrotus lividus* (*P. lividus,* Fig. 2A-F). *Hs-VEGFa(165)* overexpression results in the formation of ectopic spicule branching, 3 days post-fertilization (3dpf, (Fig. 2E,F)), similar to the phenotype of sea urchin *Pl-VEGF* overexpression (Fig. 2C,D). To verify that the observed phenotype is due to human VEGF specific-activation of the sea urchin VEGF pathway, we conducted a rescue experiment, using a sea urchin VEGF splicing morpholino antisense oligonucleotide (MO), tested before in this species (12). Embryos injected with control MO and GFP mRNA show normal skeletal rods at 2dpf while embryos injected with *Pl-VEGF* MO and GFP mRNA show severe skeletal loss and a reduction in the level of the VEGF-target, SM30 (Fig. 2G,H,K,L. VEGF control of SM30 was studied in (12, 13)). Co-injection of *Pl-VEGF* MO with either human or sea urchin mRNA partially rescues the knock-down skeletogenic phenotype and SM30 expression level, in a similar way (Fig. 2I-L). Thus, human VEGF is able to rescue sea urchin VEGF knock-down with the same efficiency as sea urchin VEGF, indicating that VEGF-VEGFR recognition is conserved despite the large evolutionary distance between these two organisms.

**Figure 2.**
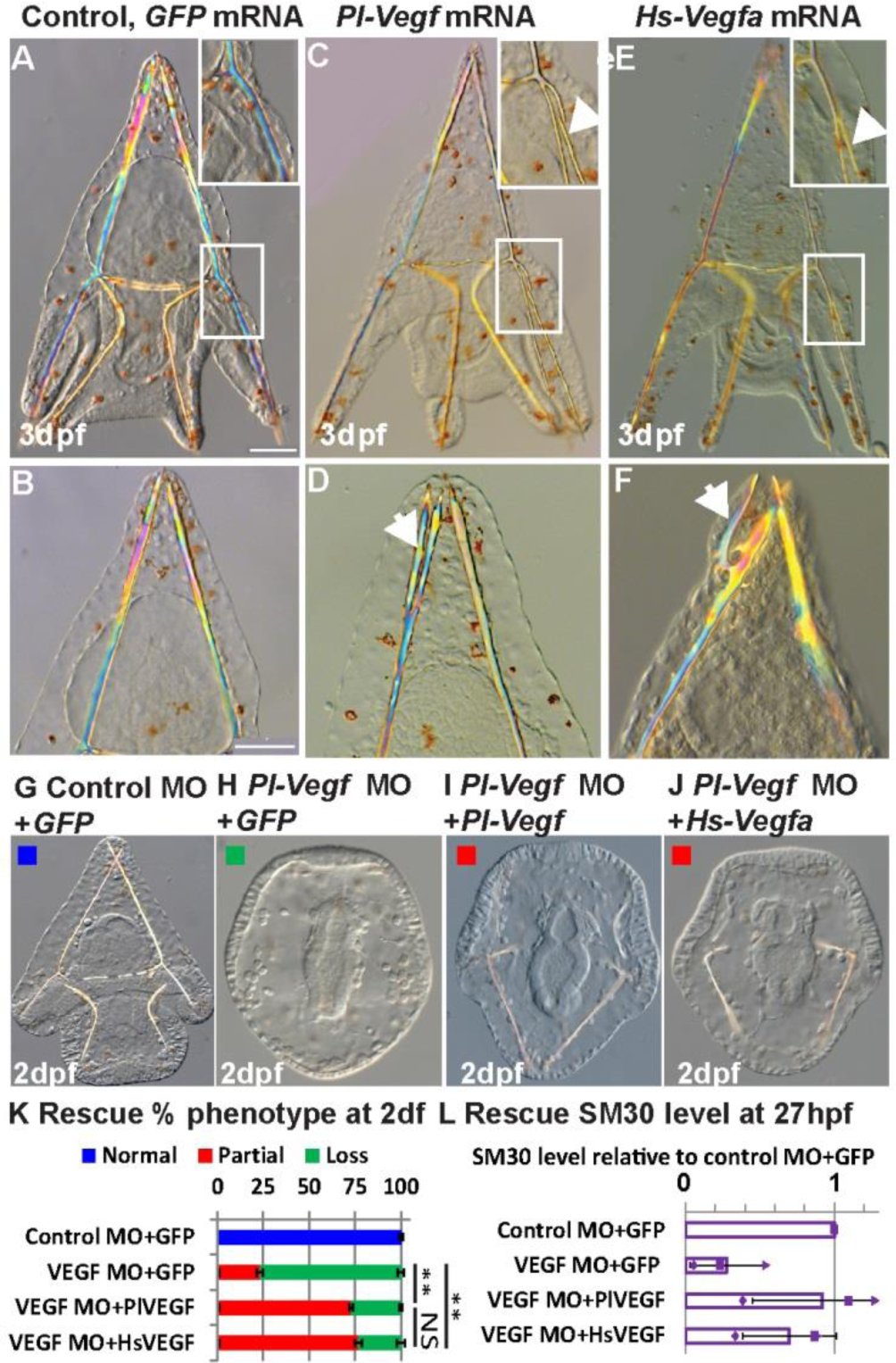
VEGF-VEGFR recognition is conserved between human and sea urchin. A-F, VEGF overexpression through 650ng/µl mRNA injection. Ectopic spicule branching on the postoral rods (top panels) and body rods (bottom panels) were observed at 3dpf in *Pl-VEGF* mRNA injected embryos (C,D; biological replicates: n=3, 52/114 embryos, 46%) and in *Hs-VEGFa(165)* mRNA injected embryos (E,F; n=4, 39/85, 46%) but not in control *GFP* injected embryos (A,B; n=4, 0/114, 0%). Scale-bars are 50µm. G-L, rescue experiment at 2dpf. G-J, embryos injected with 800mM Control MO and 650ng/µl GFP mRNA (G, normal skeleton), *Pl-VEGF* MO and GFP mRNA (H, skeletal loss); *Pl-VEGF* MO and *Pl-VEGF* mRNA (J, partial skeletal gain), *Pl-VEGF* MO and *Hs-VEGF* mRNA (J, partial skeletal gain). K, quantification of rescue phenotypes, color code is indicated in the representative images, G-J (n=3, asterisks indicates p<0.001, NS –nonsignificant, Fisher test, number of embryos scored is provided in table S1). L, SM30 mRNA levels in different treatments compared to control MO+GFP mRNA, showing average ratio (bars) and individual measurements, (QPCR). Error bars indicate standard deviation.

### VEGF is required for calcium vesicle secretion

In both tubulogenesis and biomineralization, vesicle formation and secretion play an important role. During vascular tubulogenesis, vesicles are formed in endothelial cells through pinocytosis, an apical surface is established between two adjacent cells and vesicles are secreted into the intercellular domain to form the lumen (cord hollowing (24, 25)). When VEGF signaling is inhibited, the apical surface still forms but the lumen is not generated (26). Relatedly, during sea urchin biomineralization, calcium is accumulated through endocytosis (10) and concentrated as amorphous calcium-carbonate in intracellular vesicles that are then secreted into the spicule cord where crystallization occurs (Fig. 1F, (7, 9)), however, the role of VEGF signaling in these processes has not been previously investigated.

To study the role of VEGF in calcium vesicle accumulation and secretion we compared these processes between normal embryos and embryos treated with the VEGFR inhibitor, axitinib, at different developmental stages (Fig. 3). Axitinib binds specifically to the kinase domain of human VEGF receptor (27) that is highly conserved between humans and sea urchins (Fig. S2A,B). Axitinib treatment results in analogous phenotypes to those observed in VEGF and VEGFR knock-down in *P. lividus* similar to its effect in other sea urchin species (Fig. S2C, (13)). We used calcein staining to mark calcium carrying vesicles (7) and FM4-64 to mark cell membranes in live sea urchin embryos (Fig. 3B). Calcein staining does not distinguish between the different phases of calcium within the skeletogenic cells, but can be used to track the calcium ions. Calcium vesicles are visible in all cells of the embryo, in agreement with previous studies (7, 10). VEGF inhibition does not prevent calcium vesicle accumulation in the SM cells throughout skeletogenesis (arrows in Fig. 3B), yet the calcite spicules form only in normal embryos (arrow heads in Fig. 3B). The number of calcium vesicles per area within the SM cells does not change with VEGFR inhibition before spicule formation (Fig. 3C, 16 hours post fertilization [hpf] and 20hfp, p>0.05, unpaired two tailed t-test). However, just after spicule initiation, at 24hpf, there is a significant increase in vesicle number in the SM cells in VEGFR inhibited embryos compared to control embryos, implying that vesicle secretion might be repressed by VEGFR inhibition (Fig. 3C, p=0.001, unpaired two-tailed t-test). A similar trend is observed at 30hpf but is not statistically significant (p>0.05, unpaired two tailed t-test). Overall, these observations imply that calcium carrying vesicles are present in the skeletogenic cells in VEGFR inhibition, yet the spicules do not form, possibly because vesicle secretion is inhibited. Still, the cellular and molecular processes involved in vesicle secretion downstream of VEGF signaling require further investigation.

### VEGF controls the expression of biomineralization and vascularization genes

The formation of a tubular structure and vesicle secretion in various systems require the activation of an extensive molecular tool kit that regulates cytoskeletal remodeling and other cellular mechanisms (25, 28). To identify the molecular mechanisms that sea urchin VEGF activates during spiculogenesis we explored the change in gene expression in response to VEGFR inhibition using RNA-seq before, during and after the spicules are formed (16, 20, 24 and 30 hpf, see Fig. S3A,B for experimental design). Interestingly, a major transcriptional response to VEGFR inhibition is detected only after the first stage of SM cell migration and lateral cluster formation, at the onset of spicule formation (24hpf, Fig. S3C). Relatedly, VEGF guidance of cell migration during vertebrates’ angiogenesis is mediated through direct regulation of the cytoskeleton remodeling machinery (29, 30). Possibly, similar mechanisms are activated by sea urchin VEGF signaling to guide the initial SM cell migration.

In an additional RNA-seq experiment we detected a recovery of the expression of the majority of VEGF-responsive transcripts at 24hpf and 30hpf, four and six hours after axitinib was washed, respectively (Fig. S3B,D). The observed recovery in gene expression is in agreement with the full recovery of the skeleton at the pluteus stage after axitinib wash (13). We combined the results of the time course and wash experiments at 24hpf and 30hpf to achieve high confidence in the prediction of VEGF targets (Fig. S4E, Data S1 [differentially expressed genes only] and Data S3 [quantitative RNA-seq data for all transcripts]). VEGFR inhibition significantly affects the expression of hundreds of genes at 24hpf and 30hpf, enriched with gene ontology (GO) terms related to growth factor signaling, biomineralization, cell fate specification and interestingly, vasculogenesis and circulatory system development (Fig. S4, Data S3). We studied the spatio-temporal expression and verified the response to VEGFR inhibition of a few key VEGF target genes that participate in these biological processes.

The formation of the sea urchin calcite spicules requires uptake and homeostasis of carbonate ions (31) as well as the production of spicule matrix proteins that control calcium-carbonate nucleation (32). Accordingly, within VEGF targets we observed enrichment of genes with GO terms related to carbonate homeostasis and calcium binding, such as, members of solute carrier HCO_3_^-^ transporter families (*e.g., Pl-slc26a5,* Fig. 4A), the enzyme carbonic anhydrase 7 (Fig. S5A) and different spicule matrix proteins (*e.g., Pl-SM30E*, Fig. S5B, in agreement with (12, 13)). These biomineralization genes are initially expressed broadly in the SM cells independently of VEGF signaling (20hpf, Fig. 4A, S5A,B, S6A-C). The expression of these genes then localizes to the SM cell clusters and becomes critically dependent on the VEGF pathway from 24hpf onward (Fig. 4A, S5A,B, S6A-C). This implies that the biomineralization proteins encoded by these genes could still be present in the skeletogenic cells in VEGFR inhibition at 24hpf, which precludes them from explaining the complete skeletal loss in this condition. Possibly, these biomineralization proteins are necessary but not sufficient for spicule formation or they require additional post-transcriptional activation that depends on VEGF signaling.

**Figure 4.**
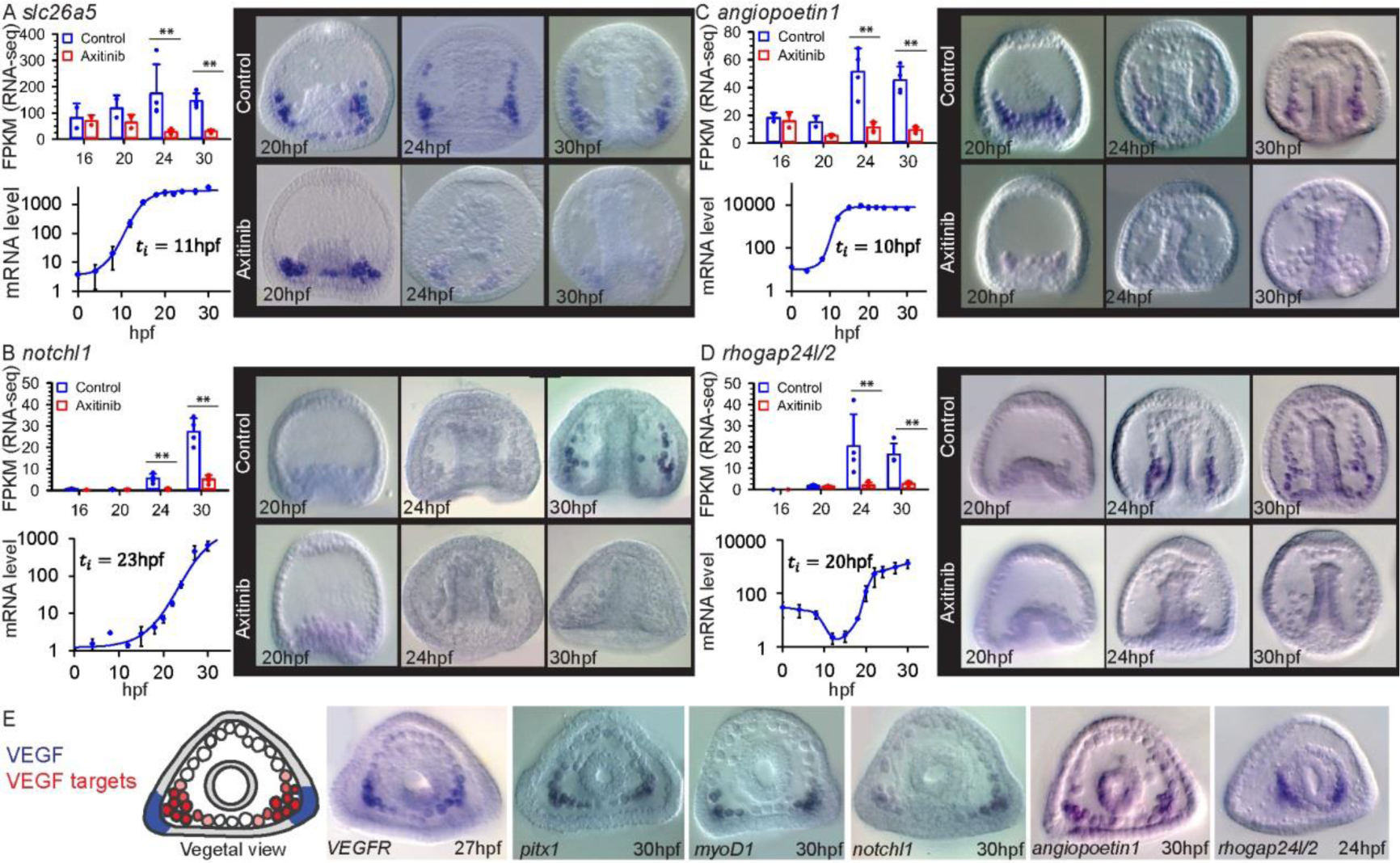
VEGF-pathway regulates the expression of biomineralization, regulatory and vascularization genes. A-D, in each panel we present: gene expression level for control and VEGFR inhibited embryos showing average expression level and individual measurements (measured by RNA-seq in Fragments Per Kilobase of transcript per Million mapped reads (FPKM), asterisks indicate FDR<0.05, 16hpf and 20hpf, n=2, 24hpf and 30hpf, n=4); temporal expression profile and initiation time (QPCR, n=3, error bars correspond to standard deviation.); spatial gene expression in control and in VEGFR inhibited embryos (whole-mount in-situ hybridization, n=3, lateral view). E, Left, illustration of *Pl-VEGF* expression (blue) and VEGF target gene expression (red). Right, examples of VEGF target gene expression in a vegetal view.

We identified several VEGF targets that have vertebrate homologs essential for vascularization or angiogenesis; *e.g., notchl1* (16), *angiopoietin1* (33) (*Sp-fred*), the cytoskeleton remodeling genes, *thsd7a* (34) (*Sp-thsd7b*)*, rhogap24l/2* (35, 36) as well as the previously reported, *VEGFR* itself (12, 13). The Notch pathway plays a prominent role in angiogenetic sprouting (16); angiopoietins are predominantly expressed at vascular supporting cells and control blood vessel number and diameter (33); human Thrombospondin type I, Thsh7A (Sp-Thsh7B) inhibits endothelial cell migration and tube formation (34). The sea urchin homologs of these genes are expressed in the skeletogenic cells and depend on VEGF activity from 24hpf and on (Fig. 4B-D, Fig. S5C,D, S6D-H). Overall, VEGF signaling becomes essential to the localized expression of both biomineralization and vascularization genes in the lateral SM clusters, which is the site of spicule formation, at the onset of spiculogenesis.

### *Pl-rhogap24l/2* is required for normal spiculogenesis

We wanted to study the role of VEGF-target, the cytoskeleton remodeling gene, *Pl-rhogap24l/2* (Fig. 4D), since cytoskeleton remodeling is critical for tubulogenesis (25) and vesicle secretion (28) in other systems. *Pl-rhogap24l/2* is homologous to the family of vertebrate genes encoding the Rho-GAPs (GTPase-activating proteins), *arhgap24, arhgap22* and *arhgap25* (35–38) (see *Pl-rhogap24l/2* phylogenetic tree, Fig. S7). Isoforms of *arhgap24* and *arhgap22* are among the highest expressed Rho-GAPs in endothelial cells (35, 36) and *arhgap25* is expressed primarily in hematopoietic cells (39). Mammalian Arhgap22, 24 and 25 are activated by RhoA, and subsequently inactivate RAC1 (37). Hence, they contribute to the counteracting interactions between RhoA and RAC1, essential for correct cytoskeleton rearrangement (36). Particularly, knockdown of *Hs-arhgap24* in Human Umbilical Vein Endothelial Cells (HUVEC) culture inhibits blood vessel formation (35).

To explore the role of *Pl-rhogap24l/2* in sea urchin skeletogenesis we studied the phenotype of *Pl-rhogap24l/2* knockdown by the injection of translation and splicing MOs (Fig. 5A-C, Fig. S8) and overexpression by the injection of mRNA of *Pl-rhogap24l/2* (Fig. 5D,E). These opposing perturbations result in an ectopic spicule branching of both the body and postoral rods at 3dpf (Fig. 5F). Possibly, the sea urchin Pl-Rhogap24l/2 contributes to the oscillations between RhoA and RAC1 activities (36, 38) and therefore either up or down-regulating it interferes with the balance between these small GTPases and impairs spicule branching. Thus, the VEGF target, *Pl-rhogap24l/2*, a sea urchin homolog of endothelial Rho-GAP genes, is activated by VEGF signaling at the SM cells and its function is necessary for normal skeletogenesis.

**Figure 5.**
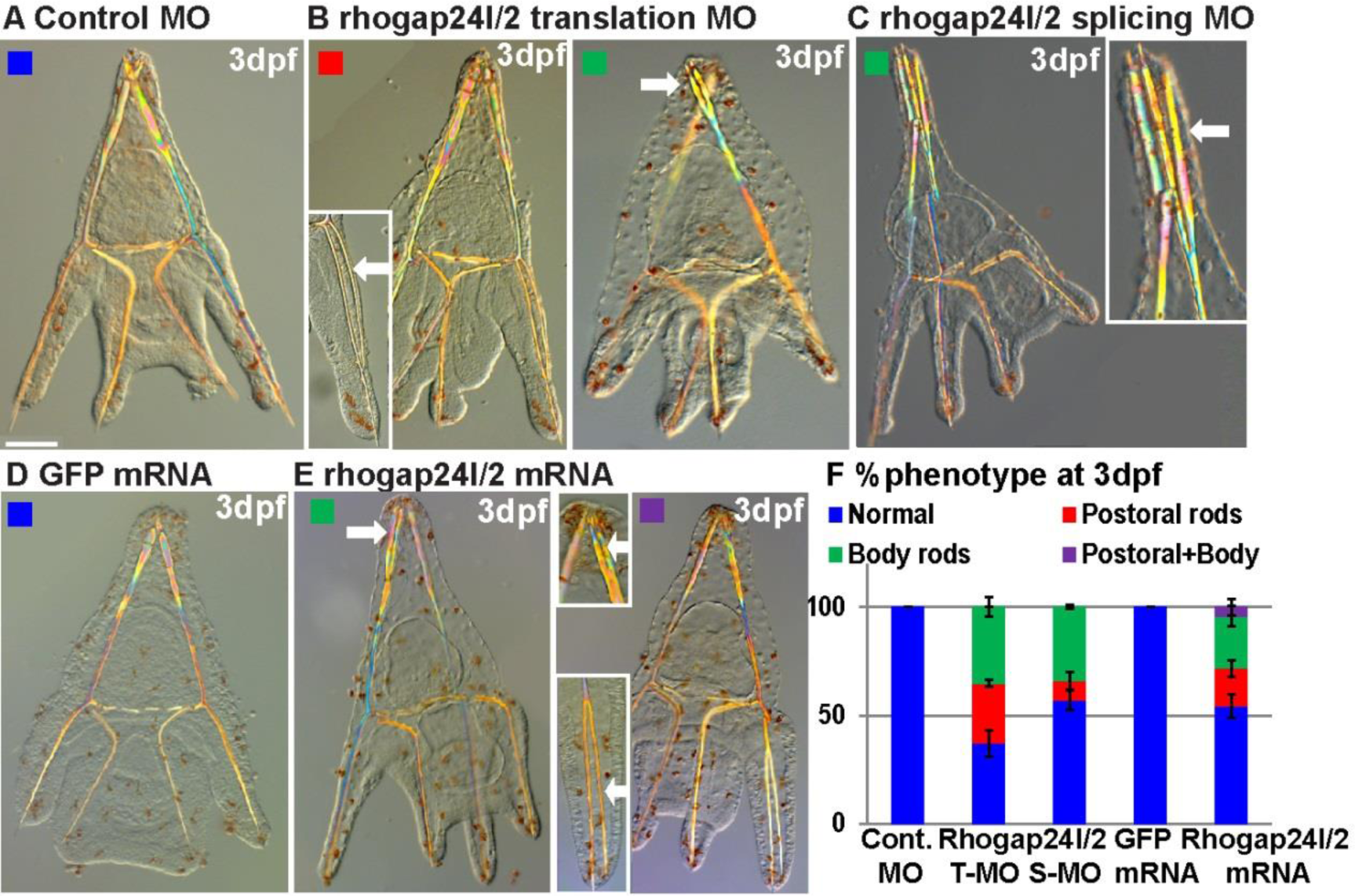
VEGF target, Pl-Rhogap24l/2, is essential for normal skeletogenesis. A-E, *Pl-rhogap24l/2* perturbations. A, embryo injected with control MO shows normal skeleton at 3dpf. *Pl-rhogap24l/*2 knock-down using either translation MO (B) or splicing MO (C) results in ectopic branching at the postoral and body rods at 3dpf. D, *GFP* overexpression results in normal skeleton while E, overexpression of *Pl-rhogap24l/2* results in ectopic branching at the postoral and body rods, sometimes within the same embryo. F, quantification of *Pl-rhogap24l/*2 perturbations, color code is indicated in the representative images A-E (n=3-5, exact number of embryos scored is provided in Table S1).

### Spiculogenesis gene regulatory network

Our results show that VEGF signaling activates its targets specifically at the lateral SM clusters located most proximally to the ectodermal VEGF secreting cells (Fig. 4E). By that, VEGF signaling could be contributing to the differential gene expression at this subset of cells that is required for spicule initiation there. Yet, VEGF is a signaling molecule and the transcriptional activation of its targets has to be mediated through a skeletogenic transcription factor. Therefore we wanted to investigate which transcription factors are expressed in the skeletogenic cell clusters at the time of spiculogenesis and whether their expression is VEGF-dependent.

The skeletogenic gene regulatory network was studied in great detail mostly at the early stages of sea urchin development (6). Some of the early skeletogenic transcription factors turn off within the skeletogenic cells hours before the spicules form (40–43) or were shown to be unaffected by VEGF perturbation (Tbr (12)). We therefore focused on the remaining six skeletogenic transcription factors: Ets1/2, Erg, Hex, Tel, FoxO and Alx1(6), each of them was shown to have a role in sea urchin skeletogenesis (6, 44). Intriguingly, vertebrate homologs of five of these transcription factors are expressed in endothelial cells and regulate different aspects of endothelial cell function (Ets1, Ets2, Erg [Erg and Fli], Hex, Tel and FOXO1 and FOX3a) (45–51).

The sea urchin genes, *ets1/2*, *erg*, *hex*, *tel*, *foxo* and *alx1* are expressed in the skeletogenic cells, including the skeletogenic clusters, at the time of spicule formation and the expression of *hex* depends on VEGF signaling (23-25hpf, Fig. 6A-F, Fig. S6K-P). Unlike the localized expression of VEGF targets, the expression of some of these transcription factors extends to the oral (*erg, tel and foxo*) and aboral (*erg, foxo and alx1*) parts of the skeletogenic ring of cells (see vegetal views in Fig. 6A-F). Of these, the most likely mediators of VEGF signaling are the ETS factors, Ets1/2, Erg and Tel of which, *ets1/2* is specifically localized at the SM lateral clusters at 24hpf (Fig. 6A). The expression of the genes *ets1/2*, *erg*, *hex* and *tel*, is detected also in non-skeletogenic mesodermal cells at the tip of the archenteron that differentiate into hematopoietic and myogenic cells (Fig. 6A-D). Thus, the combination of transcription factors expressed at the SM cell clusters at the time of spicule formation includes the VEGF-independent expression of the transcription factors, Ets1/2, Erg, Tel, FoxO, Alx1 (Fig. 6) and Tbr1 (12), as well as VEGF targets, Hex, Pitx1 and MyoD1 (Figs. 6C, S5E,F, S6I,J). We summarize our results in a partial model of the spiculogenesis GRN portraying the similarities and differences to vertebrates’ vascularization GRN (Fig. 6G).

**Figure 6.**
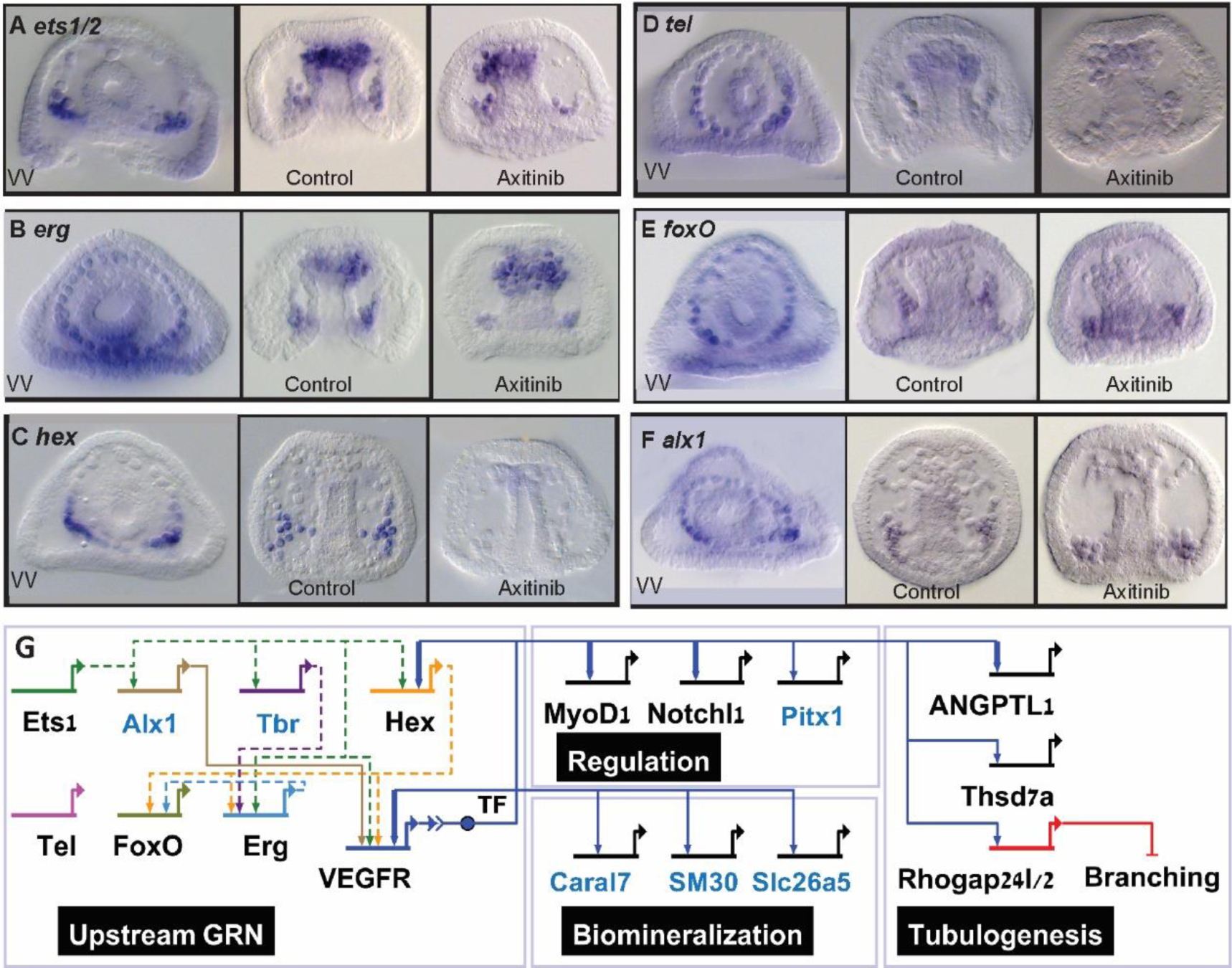
The regulatory state and VEGF downstream gene regulatory network is the skeletogenic cell clusters during the initiation of spiculogenesis. A-F Spatial expression and response to VEGFR inhibition at the initiation of spiculogenesis (23-25hpf) of the genes encoding the transcription factors, Ets1/2 (A), Erg (B), Hex (C), Tel (D), FoxO (E) and Alx1 (F), n>=3. VV-vegetal views in control embryos. G, a model of the regulatory network at the skeletogenic clusters at the time of spiculogenesis (24hpf). The regulatory interactions between the upstream transcription factors were studied only at earlier stages (6) and are therefore indicated in dashed lines. TF refers to the transcription factor that mediates VEGFR signaling. Blue gene names indicate skeletogenic specific genes and black names indicate genes common to both spiculogenesis and vascularization (46–50, 60, 67, 68). Arrows indicate activation and bars represent repression. Thick links indicate regulatory interactions observed in vertebrates’ hemangioblast differentiation (60) or in human endothelial cells (67, 68) (Data S4).

## Discussion

To gain a mechanistic understanding of the biological control of biomineralization and its rapid, parallel evolution we need to study and compare the GRNs that control it. The larval skeleton of the sea urchin is a prominent model that contributed significantly both to the understanding of GRN structure and function (6) and to the field of biomineralization (7–9). The sea urchin embryo builds its biominerals inside a tubular cavity, using VEGF signaling, a key driver of vascular systems and tubular organs in other animals (Fig. 1G). Here we unraveled some of the cellular and molecular mechanisms activated by VEGF signaling to drive sea urchin biomineralization. Our findings reveal multiple parallels to the mechanisms that drive vertebrates’ vascularization, that raise the possibility that the sea urchin biomineralization evolved through a co-option of an ancestral vascular-tubulogenesis program.

Vesicle secretion into an internal cavity is critical for both sea urchin biomineralization and for lumen formation during vascular tubulogenesis (7–9, 24). In vascularization and other systems, vesicle secretion and lumen formation are controlled by cytoskeleton remodeling proteins (25, 28). Here we show that VEGF signaling is necessary for vesicle secretion from the skeletogenic cells (Fig. 4). VEGF transcriptional target, the cytoskeleton remodeling gene, *rhogap24l/2* is necessary for normal skeletogenesis and interfering with its expression results with ectopic spicule branching (Fig. 5). Relatedly, the cytoskeleton remodeling proteins, ROCK1 and CDC42, known mediators of VEGF signaling during vascular tubulogenesis (52, 53), are critical to normal spicule formation in the sea urchin embryo (54, 55). Thus, to fully understand the molecular control of calcium vesicle secretion and spicule cavity formation it is critical to study VEGF direct control of cytoskeleton remodeling proteins.

Our studies portray the transcriptional gene network that sea urchin VEGF activates and the similarities between the spiculogenesis GRN and the GRN that controls vascularization (Fig. 6G). VEGF signaling regulates the expression of hundreds of genes at the onset of spicule formation, including regulatory, biomineralization and vascularization related genes (Fig. S3C-E, S4). The ten transcriptional targets we focused on here require VEGF signaling for their localized expression at the SM cell clusters, the subset of skeletogenic cells that generate the spicules (Fig. 4, S5). To identify the possible mediators of the transcriptional response to VEGF signaling we studied the skeletogenic regulatory state: the combination of regulatory genes co-expressed within this cell lineage (56, 57). The regulatory state is considered to be the hallmark of GRN and cell type homology that lingers throughout evolution (56, 58). Remarkably, a common set of five transcription factors (Ets1/2, Erg, Hex, Tel and FoxO) and three signaling pathways (VEGFR, Notch and Angiopoetin) essential for vascularization (16, 33, 46–50) are expressed at the sea urchin SM clusters at the time of spicule formation (Figs. 6G, black gene names). Indeed, these regulatory genes and pathways are also involved in other developmental processes across metazoans, but their co-expression in a cell lineage is unique to vertebrates’ endothelial cell and vascularization (46–50). The similarity in the downstream regulatory and effector genes is lower, but the shared downstream genes are linked to the process of tubulogenesis which is the common morphogenetic event between these different cell types. Overall, the similarity between the spiculogenesis GRN and the GRN that drives endothelial cell specification and vascularization supports a common evolutionary origin of these two networks.

Strengthening this hypothetical evolutionary scenario is the regulatory resemblance of both the sea urchin skeletogenic cells and vertebrates’ endothelial cells to the hematopoietic cell lineage and their distinction from vertebrates’ biomineralization programs. We didn’t find similarities between the sea urchin spiculogenesis GRNs and the networks that drive bone and teeth formation in vertebrates (4, 5). This is expected from the convergent evolution of biomineralization programs (2, 3). On the other hand, four transcription factors that drive sea urchin skeletogenesis and vertebrate vascularization are also drivers of hematopoietic cell differentiation, both in sea urchins (Fig. 6A-D, (59)) and in vertebrates (47, 60). Furthermore, when the skeletogenic cells are removed during sea urchin development, cells of the hematopoietic lineage change their fate and generate spicules, demonstrating the linked regulatory capacity of the two sea urchin lineages (61). The resemblance between vertebrates’ hematopoietic cells and endothelial cells is understood, as these two lineages originate from the same progenitor cells, the hemangioblasts (60). The association between the skeletogenic and hematopoietic lineages in the sea urchin could be explained if they evolved from two closely related ancestral programs of vascularization and hematopoiesis.

We speculate that the ancestral tubulogenesis program was a simple vasculogenesis program, generating a simple blood vessel, like the capillary and unlike the complex multi-layered structure of vertebrates’ arteries and veins (62). Indeed, the diameter of the spicule cord is about 4µm (Fig. 1C-E), which is of the order of the average size of the capillary (62). However, while the SM cells generate the spicule cord but keep their round mesenchymal shape (Fig. 1G), the vertebrates’ endothelial cells form an epithelial layer that constitutes the blood vessel (24, 25). Thus, there are differences not only in the filling of the tubes but also in the shape of the skeletogenic and endothelial cells that must have evolved through major changes in the ancestral GRN.

Relatedly, there are apparent differences between the spiculogenesis and the endothelial GRNs, that our model underrepresents (Fig. 6G). Multiple gene duplications have occurred in vertebrates leading to several paralogs of every sea urchin gene, for example, Ets1/2 duplication into Ets1 and Ets2, Erg duplication into Erg and Fli, and the multiple vertebrates’ paralogs of FoxO, VEGF and VEGFR (45–51). Yet, the conserved VEGF-VEGFR recognition between human and sea urchin implies that these genes originated from the same protein families and are functionally related. These gene duplications might have supported the evolution of the complex vascularization system of vertebrates with its specialized arteries and veins. Distinctive to the sea urchin skeletogenic cells are the activation of the transcription factors Tbr, Alx1 and Pitx1 and the expression of the echinoderm specific spiculo-matrix proteins and other biomineralization related genes (32) (Fig. 6G, blue genes). Furthermore, the shared transcription factors, Ets1/2, Erg, Hex, Tel and FoxO had likely acquired new targets encoding biomineralization related proteins (6). These and probably other differences between the sea urchin skeletogenic GRN and vertebrates’ vascularization GRN, had possibly contributed to the divergent outcome of these two morphogenetic programs.

In the GRN diagram we divided VEGF targets into three different functional modules: regulation, biomineralization and tubulogenesis (Fig. 6G). While in the skeletogenic cells these modules are probably tightly linked, it could be that they represent the actual building blocks from which the sea urchin skeletogenic GRN evolved. Computational simulations of the evolution of biological networks have shown that changing goals speedup evolution and leads to a modular network structure, where different blocks are responsible for different tasks (63, 64). This could be an example of the way that phylum specific biomineralization programs had rapidly evolved through the insertion of novel biomineralization modules into ancestral developmental GRNs.

## Materials and Methods

Embryos were cultured in artificial sea water (Red Sea Fish Farm LTD)) at 18°C. Exact number of biological replicates and of embryos scored for each condition in the experiments described in this work is available in Table S1.

### Imaging

Embryos were fixed and prepared for scanning electron microscopy as described in the appendix SI. Light microscopy of live embryos was done using either Ziess Axioimager M2 or Nikon A1R confocal microscope. 3D model of the spicule was generates using 52 confocal stacks by the software Imaris 7.6.5. Calcein staining was done using green calcein (C0875, Sigma, Japan) or blue calcein (Y27632, Sigma). We used FM4-64 (T13320, Life technologies, OR, USA) to stain membranes in live embryos. Further information is provided in appendix SI.

**VEGF and VEGFR protein** models were constructed based on known structures of human proteins from PDB as described in appendix SI. Protein sequence alignments were done using ClustalOmega.

### mRNA injection

cDNA of 30hpf *P. lividus* embryos was used as a template for the cloning of *Pl-VE*GF and *Pl-rhogap24l/2* cDNAs (Primer list is provided in Data S5). *Hs-VEGFa(165)* is a gift from Gera Neufeld and Ofra Kessler. mRNAs were generated and microinjected into sea urchin eggs. For VEGF rescue experiment mRNA was microinjected into sea urchin eggs along with random or VEGF splicing MO (12). Statistical analyses were done using IBM SPSS statistics version 21. Injection solutions and further details are provided in appendix SI.

**Calcium vesicle quantification** VEGFR inhibition was done using by axitinib (AG013736, Selleckchem, Houston, TX, USA), in a final concentration of 150 nM. Experiments were conducted in three biological replicates for each time point. Cell area was measured in Fiji and vesicles per cell area were counted manually by three different people. Statistical analyses were done using IBM SPSS statistics 21 as described in appendix SI.

**RNA-seq experiments** Total RNA isolation from control and VEGFR inhibited embryos was carried out using the RNeasy Mini Kit (50) from QIAGEN (#74104). Illumina libraries were constructed using NEBNext Ultra Directional RNA Library Prep Kit for Illumina (E7420). Sequencing was carried out at the Center for Genomic Technologies, The Hebrew University of Jerusalem, on an Illumina NextSeq machine (Illumina Inc., San Diego, CA) with a 100 paired-end (PE) run. Further information is provided in appendix SI. Cleaned PE reads were assembled using Trinity (version 2.0.2) PE *de-novo* assembly (65). Trinity-genes were annotated using mouse Ensembl data and the sea urchin *Strongylocentrotus purpuratus* (*S. purpuratus*) RNA-Seq assembly (http://www.echinobase.org). Trinity-genes were quantitated by EdgeR using R3.0.2. We defined significant effect of time and treatment contrasts, at P-value < 0.05 threshold, after false discovery rate (FDR) correction. Functional enrichment analysis was conducted using GOseq using *S. purpuratus* annotation (http://www.echinobase.org). Further details of the analyses are provided in appendix SI.

**Accession numbers** Raw data read sequences are available at the European Nucleotide Archive (ENA) of the EBI under accession PRJEB10269. The assembled transcriptome sequences are also available at EBI (Study PRJEB10269, accession range HACU01000001-HACU01667838).

### Rhogap24l/2 phylogenetic analysis

The 100 top blast hits for *Pl-Rhogap24l/2* were genes from the family of *rhogap22, 24* and *25* of different species. We generated a phylogenetic tree of this family using different chordates, hemichordates and echinoderm genes, as described in appendix SI (Fig. S7A). This analysis indicates that the Echinoderm predicted RhoGap24/22/25 proteins form a monophyletic clade that separated from other deuterostomes prior to paralogue formation in vertebrates.

**Quantitative polymerase chain reaction (QPCR)** was performed following the procedures outlined in (66) with some modifications described in details in appendix SI. Complete list of primer sequences is provided in Data S5.

**Whole mount in-situ hybridization (WMISH) procedure** RNA DIG probes were generated using ROCHE DIG labeling kit (catalog number 1277073910) and SP6 polymerase 10810274001 sigma. WMISH was performed as described in (43), with minor changes described in appendix SI.

**MO injection** Translation or splicing of *Pl-rhogap24l/2* was blocked by the microinjection of 800mM *Pl-rhogap24l/2* MO into sea urchin eggs (Gene Tools, Philomath, OR). Translation MO: 5-ATCCTCAAGTATCCGTAGTGTGTGA-3; Splicing MO at the 3' of the second exon: 5-TGTCCTAGAACCGTTATACTCACGT-3 (Figure S8). Injections details are provided in appendix SI.

## Acknowledgements

We thank Shlomo Ben-Tabou de-Leon and Veronica Hinman for helpful discussions. We thank Muki Shpigel and David Ben-Ezra for their help with sea urchin handling. We thank Palle von Huth for help with confocal imaging. We thank Tali Mass and Yulia Polack for help with electron microscopy. We thank Gera Neufeld and Ofra Kessler for the gift of *Hs-VEGFa(165)* plasmid. We thank Nir Sapir for his advice regarding the statistical analysis and Chuck Ettensohn, Veronica Hinman, Stefan Materna and Mark Winter for critical review of the manuscript. We thank Majed Layous, Hadeya Zaher and Nasreen Nakad for technical help with QPCR and WMISH. We thank Yarden Ben-Tabou de-Leon for the illustrations in Figs. 1G and 3A and 4E. This work was supported by the Israel Science Foundation grant number 41/14 (S.B.D.) and Eshkol post-doctoral fellowship of the Israeli Ministry of Science (M.R.).

## Author contributions

S.B.D., M.R., M.M. and T.G. designed the project. SEM embryo images were generated by T.G.. VEGF overexpression and rescue experiments were performed and analyzed by M.M.. Sequence alignment and protein structure models were constructed by M.D., T.G and S.B.D. Calcein and FM4-64 imaging were performed by M.M. with help from S.B.D. and T.G., and quantified by M.M.. M.R. designed and performed the RNA-seq experiments, N.S. prepared the libraries, M.R., and A.M. analyzed the RNA-seq data, M.R., S.B.D., T.G. and N.S. interpreted the data. T.G. and S.B.D. designed the WMISH and QPCR experiments that were performed by T.G. with help from M.M., and L.H.. M.L. made *Pl-rhogap24l/2* phylogenetic analysis. *Pl-rhogap24l/2* perturbations were designed, performed and analyzed by M.M. S.B.D. wrote the paper with significant help from M.R., T.G., M.M. and N.S..

## Author information

Raw data read sequences were deposited at the European Nucleotide Archive (ENA) of the EBI under accession PRJEB10269. The assembled transcriptome sequences are also deposited at EBI (Study PRJEB10269, accession range HACU01000001-HACU01667838). The authors declare no competing financial interests. Correspondence and request for materials should be addressed to Smadar Ben-Tabou de-Leon

