## Supplemental figures and table for "Possible co-option of a VEGF-driven tubulogenesis program for biomineralization in echinoderms"

**This PDF file includes:**

Supplementary materials and methods
Figs. S1 to S8
Table S1
Captions for databases S1 to S5
References for SI reference citations

**Other supplementary materials for this manuscript include the following:**

Datasets S1 to S5

### Supplementary Materials and Methods

**Adult animals and embryo cultures** Adult *P. lividus* were obtained from the Institute of Oceanographic and Limnological Research (IOLR) in Eilat, Israel. Spawning was induced by intracoelomic injection of 0.5M KCl. Embryos were cultured in artificial sea water (Red Sea Fish Farm LTD)) at 18°C.

**Scanning electron microcopy of sea urchin embryos** 46hpf embryos were fixed in 4% glutaraldehyde in 0.1M MOPS buffer (pH 7.0) for 24h at 4°C, then washed 3 times in MOPS buffer followed by a wash in 0.1M cacodylate buffer and post-fixed in 1% osmium tetroxide with the same buffer for 60 min at room temperature. Samples were dehydrated in an ascending ethanol series up to 100%. Samples were embedded in epon resin following manufacturer protocol (EMbed-812, Cat# 14120, Electron Microcopy Science). Thin sections (300nm) were cut and transferred to a Zeiss Sigma HD scanning electron microscope (SEM) where they were observed using a backscattered electron in-lens detector (operating at 5kV at a working distance of 4.7 mm).

**Light microscopy and imaging** All the embryos images presented in this work except from the calcein and FM4-64 staining were acquired using Ziess Axioimager M2. Calcein and FM4-64 embryo images were acquired using Nikon A1R confocal microscope. Images were later aligned in photoshop CS6 and figures were made in Adobe illustrator CS6. To generate the 3D model of the spicule, aligned fluorescent stacks from the confocal data were visualized and analyzed by Imaris 7.6.5 software. Histogram normalization was performed for stacks in which signal intensity varied greatly from slice to slice. Spicule 3D model in Fig. 1D was generated based on 50 z-stacks separated by 0.2µm of the spicule confocal images in Fig. 1C. We used the Surpass function within Imaris, in which the user indicates the channel of interest to define the specific object.

**Calcein staining** A 2mg/ml stock solution of green calcein (C0875, Sigma, Japan) or 10mg/ml stock solution of Blue calcein (Y27632, Sigma) was prepared by dissolving the chemical in distilled water. Working solution of 25µg/ml was prepared by diluting the stock solution in artificial sea water. Embryos were grown in calcein artificial sea water from fertilization and washed from calcein about two hours prior to the experiments.

**FM4-64 staining** A 100µg/ml stock solution of FM4-64 (T13320, Life technologies, OR, USA) was prepared by dissolving the chemical in distilled water. Working solution of 5µg/ml was prepared by diluting the stock solution in artificial sea water. Embryos were immersed in working solution about 10 minutes before visualization.

**VEGF and VEGFR sequence alignment and protein models** Pl-VEGF structure was modeled using PhyRe2(1) server based on Hs-VEGFD structure, PDB ID: 2xv7. This model was then superimposed on Hs-VEGFA(165) structure, PDB ID: 5HHD to get rms=0.6. Pl-VEGFR extracellular domain was modeled based on the structure of human VEGFR2-VEGFA complex, PBD ID 3V2A. This model was then superimposed on Pl-VEGF and Hs-VEGFA(165) using the structure of human VEGFR2-VEGFC complex, PDB ID: 2x1w. Pl-VEGFR kinase domain structure was modeled using Swiss-Model server(2) based on PDB ID: 1y6a. Using the structural alignment tool implemented in PyMol (https://www.pymol.org/citing) we superimposed the Pl-VEGFR-KD model with the structure of HsVEGFR2-KD-axitinib complex (PDB ID: 4ag8(3)) and obtained rms=0.715. Visualization of proteins structures was obtained using PyMol. Protein sequence alignments were done using ClustalOmega(4).

**mRNA preparation** Total RNA of 30hpf *P. lividus* embryos was used to generate cDNAs using SuperScript™ II Reverse Transcriptase (Thermo Fisher scientific 18064022). cDNAs of *Pl-VE*GF and *Pl-rhogap24l/2* were PCR amplified, ligated and inserted into PGEM plasmid (Primer list is provided in Data S5). *GFP* coding region was PCR amplified from a GFP-tag construct(5). *Hs-VEGFa(165)* is a gift from Gera Neufeld and Ofra Kessler. mRNAs were generated using Invitrogen mMESSAGE mMACHINE™ T3 Transcription Kit catalog number AM1350 and poly-adenylated using Invitrogen Poly(A) tailing kit catalog number AM1350.

**mRNA injection** mRNA was microinjected into sea urchin eggs along with 1µg/ml rhodamine dextran (D3329 Molecular probes, OR, USA) and 0.12M KCl. The injection concentration of the mRNAs was: *Pl-VEGF*, *Hs-VEGFa(165)* and *GFP* mRNA 650 ng/µl (Fig. 2), *GFP* and *Pl-rhogap24l/2* mRNA 800 ng/µl (Fig. 5D, E). Exact number of biological replicates and of embryos scored for each condition is available in T S1. Summary of the phenotypes for each experiment is provided in the paper's text and figures.

**VEGF rescue experiment** mRNA in the concentration indicated above was microinjected into sea urchin eggs along with 800 µM of random or VEGF splicing MO(6), 1µg/ml rhodamine dextran (D3329 Molecular probes, OR, USA) and 0.12M

KCl. Exact number of biological replicates and of embryos scored for each condition is available in extended data table 1. Summary of the phenotypes for each experiment is provided in the paper's text and figures.

Statistical analyses were done using IBM SPSS statistics version 21. Pearson's two-sided chi-square was used in intergroup comparisons of 2×2 categorical variables expressed as numbers. The chi-square test was performed to assess the association between the different treatments to the observed phenotypes (partial skeleton or skeletal loss), generating the following results:

| **Intergroup comparison** | **$\chi^2$ (df=1)** | **P-value** |
|---|---|---|
| *Pl-VEGF* MO + GFP mRNA<br>*Pl-VEGF* MO + *Pl-VEGF* mRNA | 60 | <0.0001 |
| *Pl-VEGF* MO + GFP mRNA<br>*Pl-VEGF* MO + *Hs-VEGFa(165)* mRNA | 73 | <0.0001 |
| *Pl-VEGF* MO + *Pl-VEGF* mRNA<br>*Pl-VEGF* MO + *Hs-VEGFa(165)* mRNA | 0.364 | 0.546 |

**Axitinib (AG013736) treatment** Axitinib binds specifically to the kinase domain of human VEGF receptor(7, 8) that is highly conserved between human and sea urchin (Extended data Fig. 3a,b). Axitinib treatment results with analogous phenotypes to those observed in VEGF and VEGFR knock-down in *P. lividus* (Fig. S3C)(6), similarly to its effect in other sea urchin species(9). A 5mM stock solution of the VEGFR inhibitor, axitinib (AG013736, Selleckchem, Houston, TX, USA), was prepared by reconstituting this chemical in dimethylsulfoxyde (DMSO). Treatments were carried out by diluting aliquots of the axitinib stock in embryo cultures to provide a final concentration of 150 nM. Control embryos in all experiments were cultured in equivalent concentrations of DMSO at no more than 0.1% (v/v). It was determined in a set of preliminary dose-response trials that at 150nM of axitinib 100% of *P. lividus* embryos were overall healthy displayed loss of skeleton, consistent with previous reports of VEGF morphants at 48hpf (6, 9).

**Calcium vesicle quantification** Experiments were conducted in three biological replicates (separate pairs of parents) for each time point. In each biological replicate, vesicle number in at least eight cells from three different embryos was measured for each condition resulting in a range of 53-71 cells for each condition. Cell area was measured in Fiji and vesicles per cell area were counted manually by three different people. Statistical analyses were done using IBM SPSS statistics 21. Continuous outcome variables (#vesicle/$\mu m^2$) exhibiting a skewed distribution were transformed, using the natural logarithms (16 and 20hpf) or square roots (24 and 30hpf) before t-tests were conducted to satisfy the prerequisite assumptions of normality by Kolmogorov-Smirnov test. A Student unpaired, two-tailed t-test was used for comparison of between-group data. P-values $< 0.05$ were considered statistically significant. An independent samples t-test was performed comparing the mean consistency scores of #vesicle/$\mu m^2$ in skeletogenic cells, in control and in VEGFR inhibited embryos generating the following results, 16hpf: Control (M=0.04, SD=0.02); VEGFR inhibition (M=0.04, SD=0.02); t(132)=-0.578, P=0.564). 20hpf: control (M=0.06, SD=0.03); VEGFR inhibition (M=0.05, SD=0.03); t(124)=1.208, P=0.229). **24hpf**: control (M=0.06, SD=0.02); VEGFR inhibition (Mean, M=0.08, SD=0.04); t(123.691)=-3.34, **P=0.001**). 30hpf: control (M=0.06, SD=0.02); VEGFR inhibition (M=0.07, SD=0.04); t(115.704)=-1.343, P=0.182). Here M indicates mean, SD is standard deviation, t(df)=t-value, P is P-value. Mann-Whitney Nonparametric analyses produced similar results (16hpf p=0.661, 20hpf p=0.112, **24hpf p=0.002**, 30hpf p=0.463). Graphs in Fig. 3C (box plots) were generated using SigmaPlot software.

**RNA-seq experimental design and sampling procedure** The effects of VEGF inhibition on gene expression in sea urchin embryos was assessed in two separate but related experiments 1) a 'time-course' experiment in which embryos were cultured continuously in the presence of axitinib prior to sampling at 16, 20, 24 or 30hpf, and 2) a 'wash' experiment in which embryos were cultured either continuously with axitinib or until it was washed out at 20 or 24hpf, prior to sampling at 24 or 30hpf, respectively (Fig. S3B). Each of these two experiments consisted of eggs and sperm from two independent pairs of parents (N=4 total). Total RNA isolation from control and treated embryos was carried out using the RNeasy Mini Kit (50) from QIAGEN (#74104) according to the kit protocol. We confirmed axitinib affect in representative embryos from all experimental sets used for RNA-Seq analyses by visually inspecting a skeleton-loss at 48hpf.

**RNA-seq sample preparation and sequencing** Extracted RNA was treated with TURBO DNase (Ambion, AB-AM1907, Austin, TX, USA) according to the manufacturers' instructions and assessed using capillary electrophoresis (Agilent Bioanalyzer 2100, Agilent Technologies, Santa Clara, CA). All samples had a RIN of 8 or higher. Illumina libraries were constructed for all samples (n=28) using NEBNext Ultra Directional RNA Library Prep Kit for Illumina (E7420) with 1µg starting material according to the manufacturer's instructions. Sequencing was carried out at the Center for Genomic Technologies, The Hebrew University of Jerusalem, on an Illumina NextSeq machine (Illumina Inc., San Diego, CA) with a 100 paired-end (PE) run with all libraries multiplexed together on all lanes to remove the possibility of batch effects. Approximately 30 million reads were obtained per sample.

**Transcriptome assembly and annotation** PE reads were cleaned from adapters, and low-quality regions, using Trimmomatic 0.3. The quality of reads was further inspected using Fastqc (http://www.bioinformatics.babraham.ac.uk/projects/fastqc/). PE reads were assembled using Trinity (version 2.0.2) PE de-novo assembly (10, 11), with strand specific RF library (SS_lib_type RF option). Totally, 667,838 contigs were assembled in 490,852 Trinity-gene-groups (corresponding to putative isoforms vs. genes or genes segments). We compared the obtained *P. lividus* contigs to protein predictions of the sea urchin *Strongylocentrotus purpuratus* (*S. purpuratus*) genome-based RNA-Seq assembly (http://www.echinobase.org)(12), using blastx. For each query contig, the top blastx *S. purpuratus* hit was selected (if available), after filtering out hits with e-value>$10^{-5}$. In total, 165,175 *P. lividus* contigs, belonging to 115,798 Trinity-genes, were annotated using *S. purpuratus* data. Similarly, 101,003 *P. lividus* contigs, belonging to 69,831 Trinity-genes were annotated using mouse Ensembl data. Raw data read sequences are available at the European Nucleotide Archive (ENA) of the EBI under accession PRJEB10269. The assembled transcriptome sequences are also available at EBI (Study PRJEB10269, accession range HACU01000001-HACU01667838).

**RNA-seq gene-level expression quantitation and differential expression analysis**

*P. lividus* PE reads were mapped to all available contigs, via RSEM 1.2.6(13), using the Trinity information of genes/isoforms hierarchy. We further calculated expression levels and differential expression for time course and wash experiments separately, and for merged data from both experiments (provided as different pages in Data S1). For the foregoing three analyses (time course, wash and merged data), 49,627, 48,644, and 53,596 Trinity-genes with Counts Per Million mapped reads (CPM) > 3 in at least one sample, were quantitated by EdgeR using R3.0.2 (14). From these groups, 19,082, 18,750, and 22,002 were mapped to *S. purpuratus* genes, respectively. In EdgeR, trimmed mean of M values (TMM) normalization was first conducted(15), and then CPM and FPKM (Fragments Per Kilobase of transcript per Million mapped reads) values were obtained. Differential expression analysis was conducted using Edger (R 3.0.2). ). In order to detect the foregoing batch, treatment and time effects, at the gene level, we used blocking design in the EdgeR GLM framework. We defined significant effect of time and treatment contrasts, at P-value < 0.05 threshold, after false discovery rate (FDR) correction (16).

**Gene Ontology functional enrichment analysis** Functional enrichment analysis was conducted using GOseq(17). A custom GOseq GO database was built using the publically available Blast2Go *S. purpuratus* annotation for WHL22 version (http://www.echinobase.org). Specifically, the level of enrichment was calculated by comparing the proportion of genes belonging to specific functional categories among differentially-expressed (DE) genes (foreground), compared to their proportion among all expressed genes (background). Enrichment significance was defined at P-value < 0.05 threshold, after FDR correction.

**Homology and phylogenetic Analyses of Pl-Rhogap24l/2** The top 100 blast hits for Pl-Rhogap24l/2 were proteins encoding Rho-GAPs 24, 22, and 25 in different species, *e.g. S. purpuratus*, *Sp-rhogap24l/2* (evalue=0, identity 90%) and mouse Mm-Argpag24 (evalue=$10^{-63}$, identity 37%, Data S2). Vertebrates' Arhgap22, 24 and 25 proteins form a family of structurally similar Rho-GAPs that are expressed in different tissues and related to various functions (18-21). To study the orthology relationship between echinoderm and vertebrates Rhogap24/22/25 we generated a phylogenetic tree of these genes: We first aligned human RhoGap22, RhoGap24, and RhoGap25 sequences (NP_001020787, NP_001242954 and NP_055697, respectively) against Blast NR database, selecting the highest homologies among different Chordates (Vertebrates, Tunicates, Cephalochordates), Hemichordates and Echinoderm species. Amino acid sequences were aligned using MAFFT V.7.036 (http://mafft.cbrc.jp/alignment/server/) (22). Multiple sequence alignments were manually reviewed for minor corrections. We then performed Bayesian inference of phylogeny of these proteins, using gamma distributed rate variation across sites, as implemented in MrBayes (23). MrBayes analysis was run for 1,000,000 Markov chain Monte Carlo generations, sampling trees every 5,000 generations and discarding the first 25% of samples as the burn-in fraction, as suggested by the authors (23). Two Bayesian chains were run to ensure adequate mixing. Convergence was indicated by an average standard deviation of split frequencies (ASDSF) <0.01 between the two chains and a potential scale reduction factor (PSRF) value ~1 for all parameters. The 50% majority consensus tree was selected as the final tree. This analysis indicates that the Echinoderm predicted RhoGap24/22/25 proteins form a monophyletic clade that separated from other deuterostomes prior to paralogue formation in vertebrates (Fig. S7). Specifically, Pl-Rhogap24l/2 does not have a true one-to-one vertebrate ortholog but is one of the homologs to the vertebrates' RhoGap24/22/25 proteins.

**Quantitative polymerase chain reaction (QPCR)** QPCR was performed following the procedures outlined in(24) with some modifications:

***cDNA preparation for QPCR*** Rescue experiments: Total RNA was extracted from 200 sea urchin embryos in each condition using RNeasy Micro Kit (50) from QIAGEN (#74004) according to the kit protocol using DNase treatment from RNease-Free DNase Set- Qiagen (50) (#79254). The total RNAs were reverse-transcribed using the High Capacity cDNA RT kit, AB-4368814 (Applied Biosystems) following the manufacturer's instructions.

VEGFR inhibition: Total RNA was extracted from >1000 sea urchin embryos in each condition using RNeasy Mini Kit (50) from QIAGEN (#74104) according to the kit protocol using DNase treatment from RNease-Free DNase Set- Qiagen (50) (#79254).

1µg of total RNAs were reverse-transcribed using the High Capacity cDNA RT kit, AB-4368814 (Applied Biosystems) following the manufacturer's instructions.

Subsequently, QPCR was carried out in triplicates using a 384CFX-real time thermal cycler and the IQ SYBR Green Supermix (Bio-Rad laboratories, Hercules, CA). Reaction conditions were as follows: 95°C for 3 min (one cycle), followed by 95°C for 10 s, 55°C for 30s (40 cycles). Dissociation analysis was performed at the end of each reaction to confirm the amplification specificity. Primer sets for all tested genes were designed using Primer3Plus (http://www.bioinformatics.nl/cgi-bin/primer3plus/primer3plus.cgi/) and a complete list of their sequences is provided in Data S5.

***QPCR expression level quantification and differential expression analysis***

Time course (Fig. 4 and Figs. S3, S6): To quantify the relative levels of mRNA per sample we inserted a known number of GFP cDNA molecules to each sample that includes cDNA transcribed from 1.25ng of extracted total RNA of *P. lividus* at different time points as described in(24). Every experimental time point for each gene was replicated in three biological replicates (three sets of parents) each sample was assessed in three technical replicates. The calculation of gene prevalence compared to GFP was performed using the formula, $\mathrm{GFP} \times 1.9^{(CtGFP-Ctgene)}$, with a constant coefficient efficiency factor, 1.9, corresponding to the average value of all primers set. Gene initiation times were calculated using a sigmoid fit: $\log(mRNA(t)) = a - b/(1 + \exp(c(t - \boldsymbol{ti}))$, considering $t_i$ as the estimated initiation time (24, 25). To fit we used Matlab's Curve Fitting Toolbox, using the nonlinear least-squares method.

Differential expression in rescue experiments and in VEGFR inhibition and (Figs. 2, Figs. S2, S7): For measuring changes in genes expression at different condition at the same time point, the relative expression of genes was determined by $1.9^{-\Delta\Delta CT}$ method using Ubiquitin as an internal gene reference for normalization (26). Then the ratio between gene-expression levels in different treatments and in control embryos was calculated for each biological replicate and time point. Every experimental time point for each gene was replicated in three biological replicates (three sets of parents) each sample was assessed in three technical replicates.

**Whole mount in-situ hybridization (WMISH) probe preparation** Total RNA of 30hpf *P. lividus* embryos was used to generate cDNA using the High capacity cDNA RT AB-4368814 (Applied Biosystems). cDNAs of the tested genes were PCR amplified, ligated and inserted into pGemT (Promega A3600) or pJet plasmids (Thermofisher scientific K1231). Primer list is provided in Data S5. RNA DIG probes were generated using ROCHE DIG labeling kit (catalog number 1277073910) and SP6 polymerase 10810274001 sigma.

**WMISH procedure** WMISH was performed as described in (27), with minor changes. Embryos were fixed in 4% paraformaldehyde, 33mM Maleic acid buffer pH7, 166mM NaCl, overnight at 4˚C. The embryos were rehydrated by five washes with MABT buffer. Hybridization was conducted overnight at 60-65˚C in 50% formamide, 5XSSC, 5xDendards (sigma D2532), 1mg/ml yeast tRNA, 5mg/ml Heparin, 0.1% Tween-20. After washes with SSC, embryos were incubated in DIG epitope detection buffer: 0.1M Maleic acif pH7, 0.5M NaCl, 10% sheep serum, 0.1mg/ml BSA, 0.1% Tween-20. The staining reaction was done in the AP buffer containing 10% dimethylformamide and NBT/BCIP. Each probe was studied in at least 2 biological replicates where at least 30 embryos were observed for each condition.

---

**Rhogap24l/2 MO injection** Translation or splicing of *Pl-rhogap24l/2* was blocked by the microinjection of *Pl-rhogap24l/2* MO into sea urchin eggs. MO was synthesized (Gene Tools, Philomath, OR) complementary to the sequence of *Pl-rhogap24l/2*. Translation MO: 5-ATCCTCAAGTATCCGTAGTGTGTGA-3; Splicing MO at the 3' of the second exon: 5- TGTCCTAGAACCGTTATACTCACGT-3. The injection of the splicing MO generated a deletion of the second half of the second exon that contains the PH domain, as tested by PCR of cDNA prepared from injected embryos (Figure S8, PCR kit: FIREPol master mix 042500115 solisbiodyne, primers are provided in Data S5). Microinjection solutions were prepared containing 800mM *Pl-rhogap24l/2* MO, 0.12 M KCl and 1µg/ml Rhodamine Dextran. Embryos injected with similar concentration of random MO were used as a control. Three biological replicates were studied at 72hpf (see table S1 for exact numbers of embryos scored in each condition).

### Supplementary figures and tables

A
```
Hs-VEGFa(165)      MNFLLSWVHWSLALLLYL------------------------------------------
Pl-VEGF            -MYMGNIKPWTLSFIFYLFVIVLSHQVESTHSVPMLSQRVIEESMSNVNSLEEYLSALSL
                     :: .    *:*::::**

Hs-VEGFa(165)      -------------------HHAKWSQAAPM-AEGGGQNHHEVVKFMDVYQRSYCHPIETL
Pl-VEGF            NGTSTLSSSYYLNSSPTALNKSSYSSLGRAGSAGGLQNSAFLASLEDENARVQCQPRDRV
                                      :::.:*. .    : ** **    :..: *    *   *:* : :

Hs-VEGFa(165)      VDIFQEY--PDEIEYIFKPSCVPLMRC--GGCCNDEGLECVPTEESNITMQIMRIKPHQG
Pl-VEGF            VDSYEELGIPSGYGDFLYPECIVVRRCKQGGCCGDE-SECVPSSTTNITMNFLKTGSEIT
                   ** ::*    *.     :: *.*: : **  ****.**   ****:. :****:::    .

Hs-VEGFa(165)      QHIGEMSFLQHNKCECRPKKDR--------------ARQENPCGPCSERRKHLFVQDPQT
Pl-VEGF            REI-----VNDLECECQDNPSFCPEPVVDCPNNKVWSYSE-CTCKCRNRCPKPFLQDEDT
                   :.*      ::. :***: :  .                  : .*      * :*   : *:** :*

Hs-VEGFa(165)      CKCSCKNTDSRCKARQL-----ELNERTCRCDKPRR------------------------
Pl-VEGF            CGCDCLSQDSNCKNIYSGSRNGKLSREECDCVRNGLCGYPPCINGGFSISDCNCINSNSW
                   * *.* . **.**         :*... * * :
```

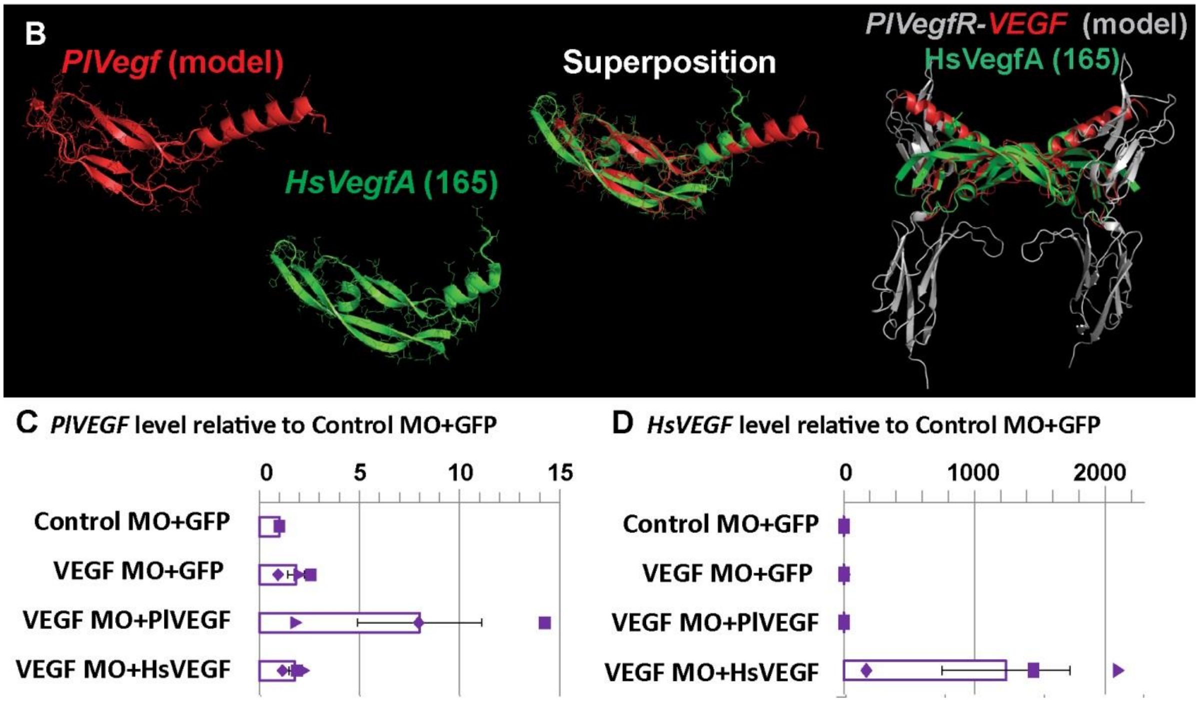


**Figure S1 Human vs. sea urchin VEGF sequences and rescue.** A, protein sequence alignment between the sea urchin *Pl-VEGF* and human *Hs-VEGFa(165)*. Asterisk indicates amino acid identity, highly similar amino acids are indicated in two dots and similar amino acids are indicated in one dot. B, Left, a superposition of the model of sea urchin Pl-VEGF structure (red, based on PDB ID: 2xv7) and human VEGFa(165) isoform (green, PDB ID: 5hhd) shows similarity between the structures (rms=0.6). Right, Superposition of VEGFa(165) isoform on a model of Pl-VEGF-PlVEGFR complex (red and gray, based on PBD ID 3V2A). C,D, mRNA levels in different treatments compared to control MO+GFP mRNA at 27hpf (QPCR). C, *Pl-VEGF* level increases when exogenous *Pl-VEGF* is injected; D, *Hs-VEGFa(165)* mRNA is detected by QPCR in embryos injected with *Hs-VEGFa(165)*. Bars show averages, markers indicate individual measurement, error bars indicate standard deviation.

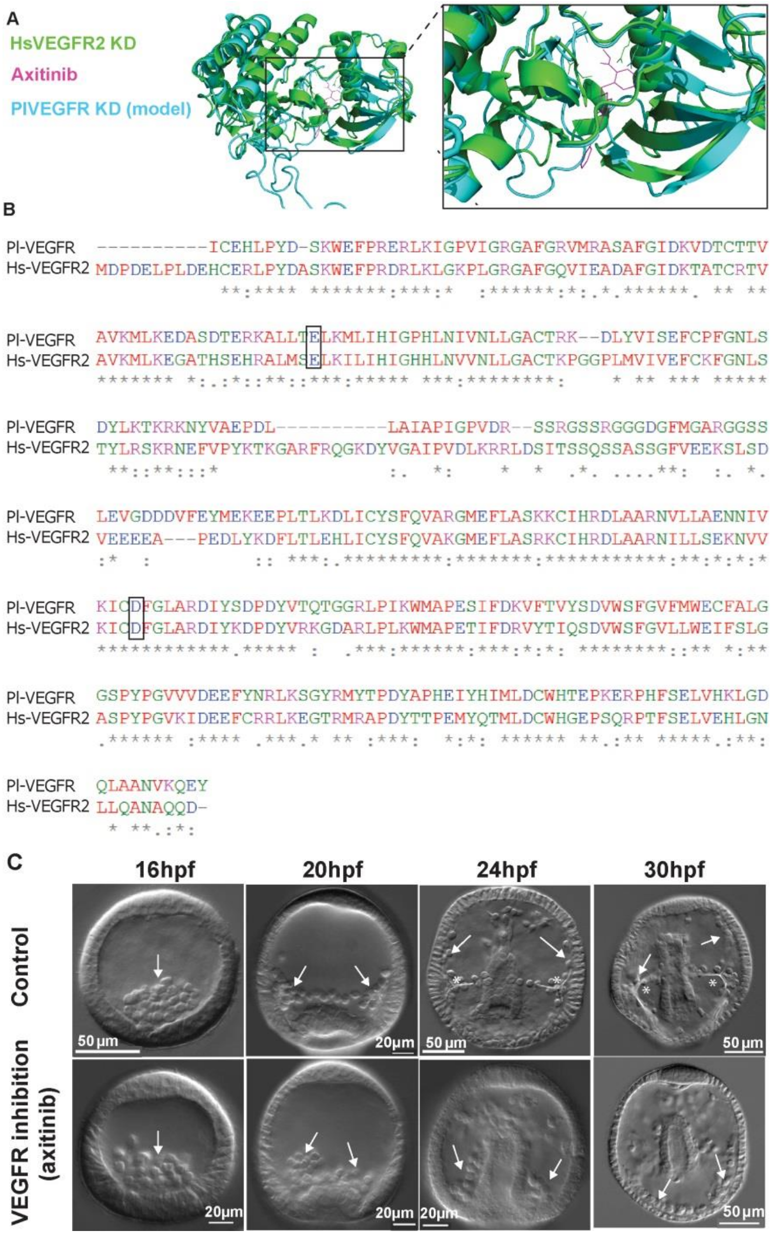


**Figure S2 A comparison of VEGFR kinase domain and axitinib binding between human and sea urchin and axitinib phenotypes in sea urchin.** A, high similarity between sea urchin VEGFR predicted structure (model based on PDB ID: 1y6a) and the structure of VEGFR2-axitinib complex (PDB ID: 4ag8(3), rms=0.715). B, protein sequence alignment of sea urchin VEGFR and human VEGFR2 kinase domain. Rectangles mark the amino acids where axitinib binds(3, 28). Asterisk indicates amino acid identity, highly similar amino acids are indicated in two dots and similar amino acids are indicated in one dot. C, phenotype of VEGFR inhibition using axitinib are similar to VEGF and VEGFR knock-down experiments in *P. lividus*[9] and in other sea urchin species[10]. White arrows point to the SM cells, asterisks denote the spicules. Epithelial to mesenchymal transition happens in both control and VEGFR inhibited embryos (16hpf), yet the distribution of the SM cells is distorted and the lateral clusters are not formed (20hpf). At 24hpf and 30hpf the spicules form and elongate in control embryos and some of the SM cells migrate toward the animal pole; these processes do not occur in VEGFR inhibition[9,10].

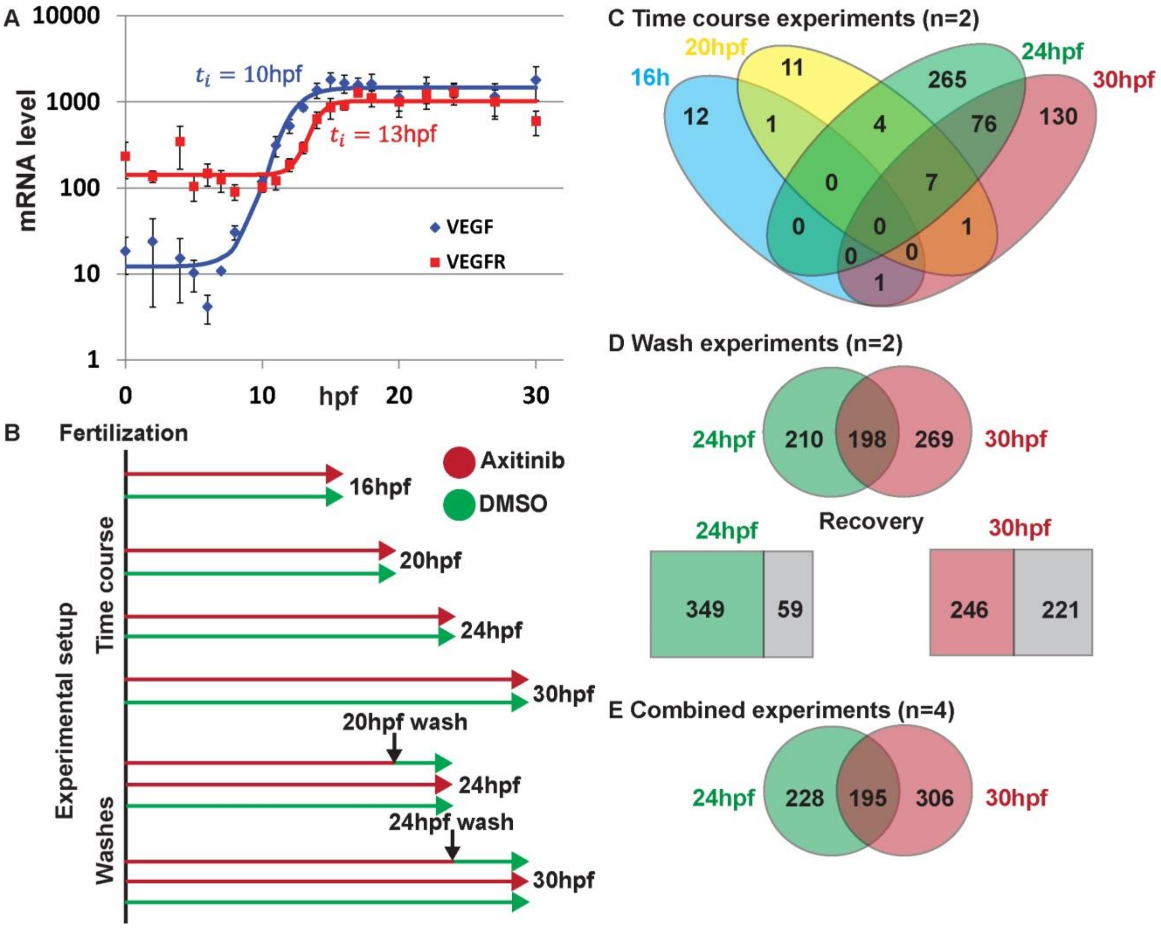


**Figure S3 RNA-seq experimental design and results.** A, *Pl-VEGF* expression turns on at 10hpf and *Pl-VEGFR* expression is activated at 13hpf (QPCR). B, he design of the RNA-seq experiment. Embryos were grown in artificial sea water with 150nM of VEGFR inhibitor, axitinib, and with DMSO as control. Embryos were collected for RNA extraction at the times depicted in the figure. In the wash experiments, a third culture of embryos was grown in axitinib and washed at the time indicated in the figure. C. number of genes that were differentially expressed in VEGFR inhibition compared to control at the four experimental time points (FDR<0.05). D, number of genes that were differentially regulated in VEGFR inhibition at 24hpf and 30hpf in the wash experiments (circles, FDR<0.05). Portion of the genes that recovered after axitinib wash at 24hpf (349/408, 86%) and at 30hpf (246/467, 53%). E, differential gene expression based on the time course and wash experiments combined at 24hpf and at 30hpf showing the overlap between differentially expressed genes at 24hpf and 30hpf (FDR<0.05).

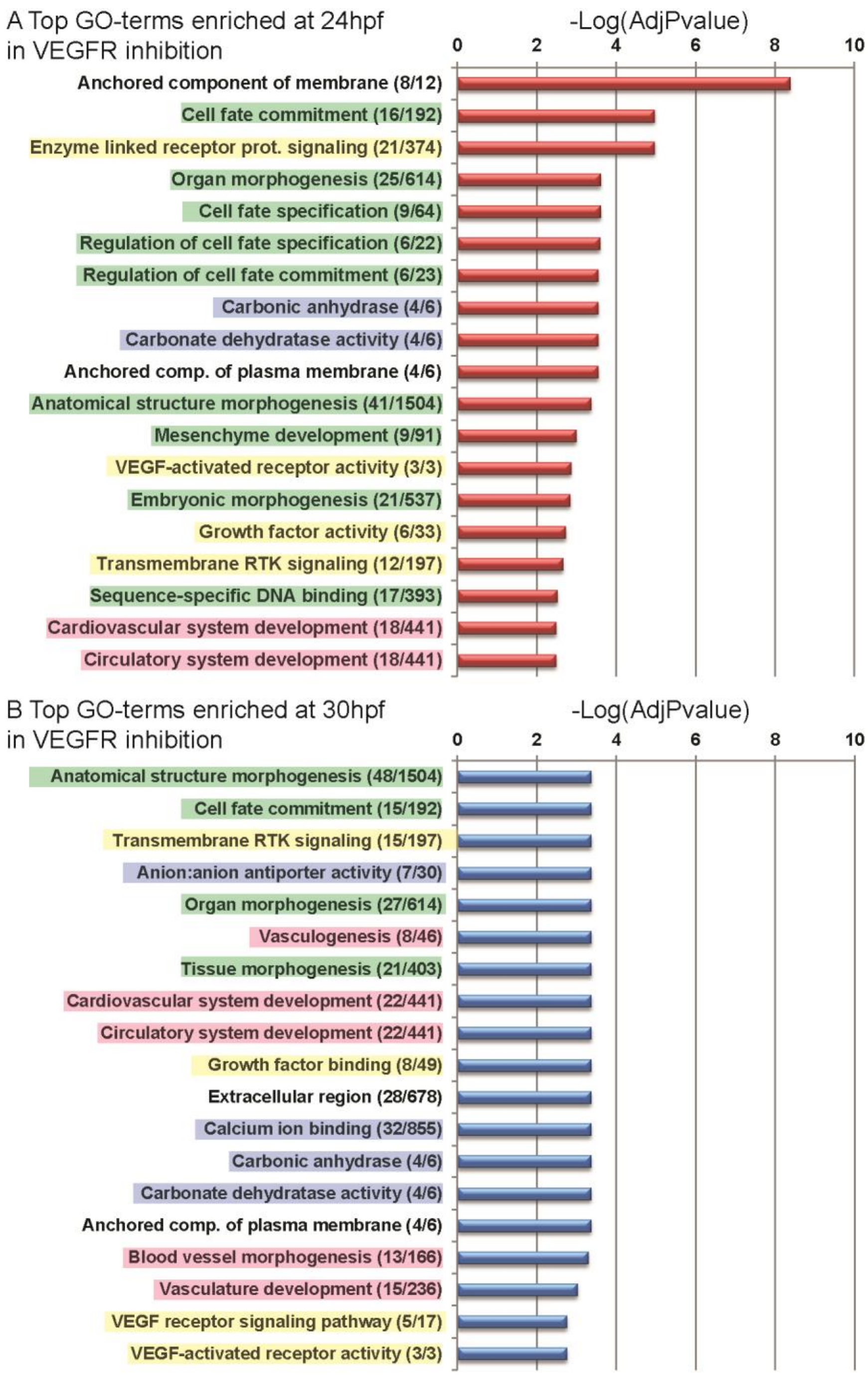


**Figure S4 GO-enrichment of differentially expressed genes under VEGFR inhibition at 24hpf and 30hpf.** Overrepresentation at 24hpf (A) and at 30hpf (B) was calculated using GO-seq and values are presented as –Log(Adjusted P-value), see methods for details. Numbers in the brackets indicate the number of genes with this GO-term within VEGFR targets vs. the background (all transcripts). Color highlights GO-terms related to the following biological processes: green – embryo development and cell fate specification, yellow – growth factor signaling, blue – biomineralization, pink – vascularization.

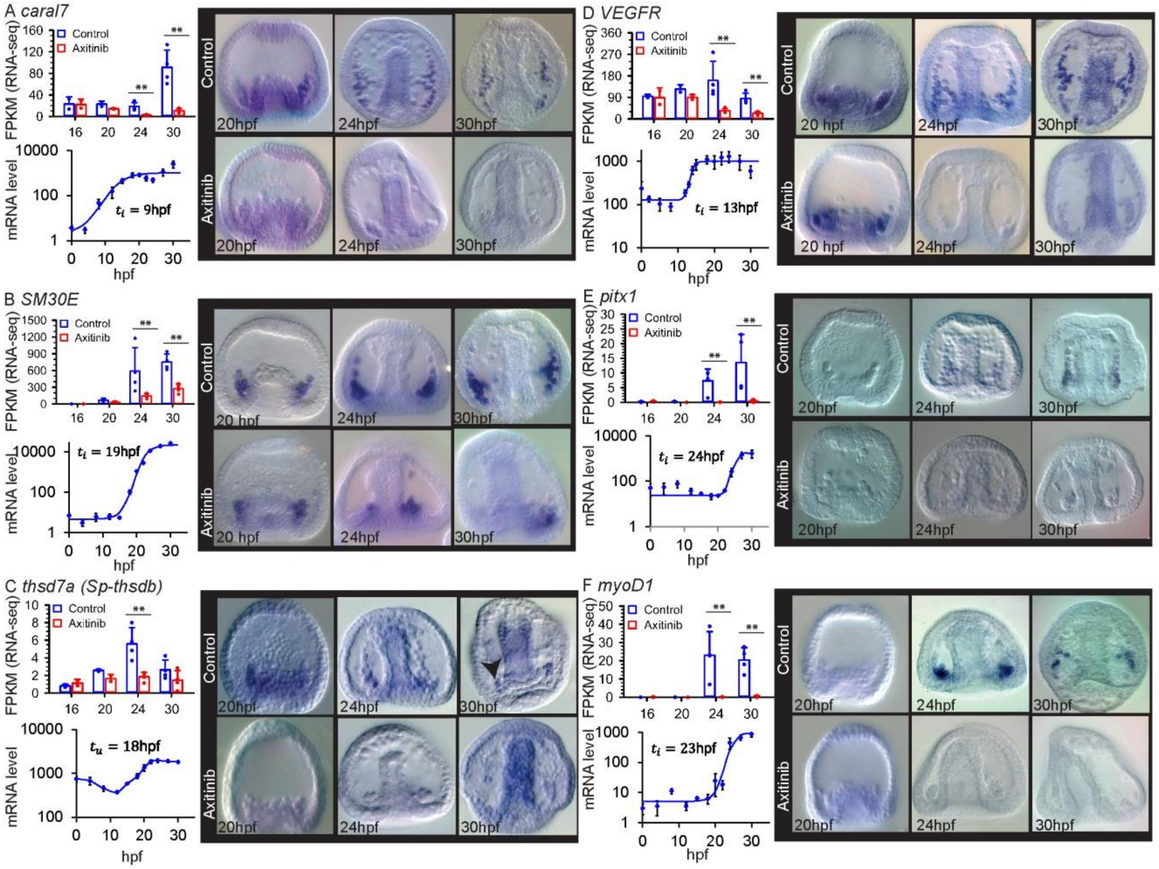


**Figure S5 VEGF signaling controls the expression of regulatory, biomineralization, and vascularization related genes.** In each panel in we present: gene expression level according to RNA-seq results for control and VEGFR inhibited embryos (measured in Fragments Per Kilobase of transcript per Million mapped reads (FPKM), 16hpf and 20hpf, n=2, 24hpf and 30hpf, n=4); gene temporal expression profile and initiation time measured by QPCR (n=3); the spatial expression of the gene in control embryos and VEGFR inhibited embryos at three developmental time points (whole-mount in-situ hybridization, vegetal view). Error bars correspond to standard deviation. Dots in RNA-seq are individual data points and asterisks indicate FDR<0.05.

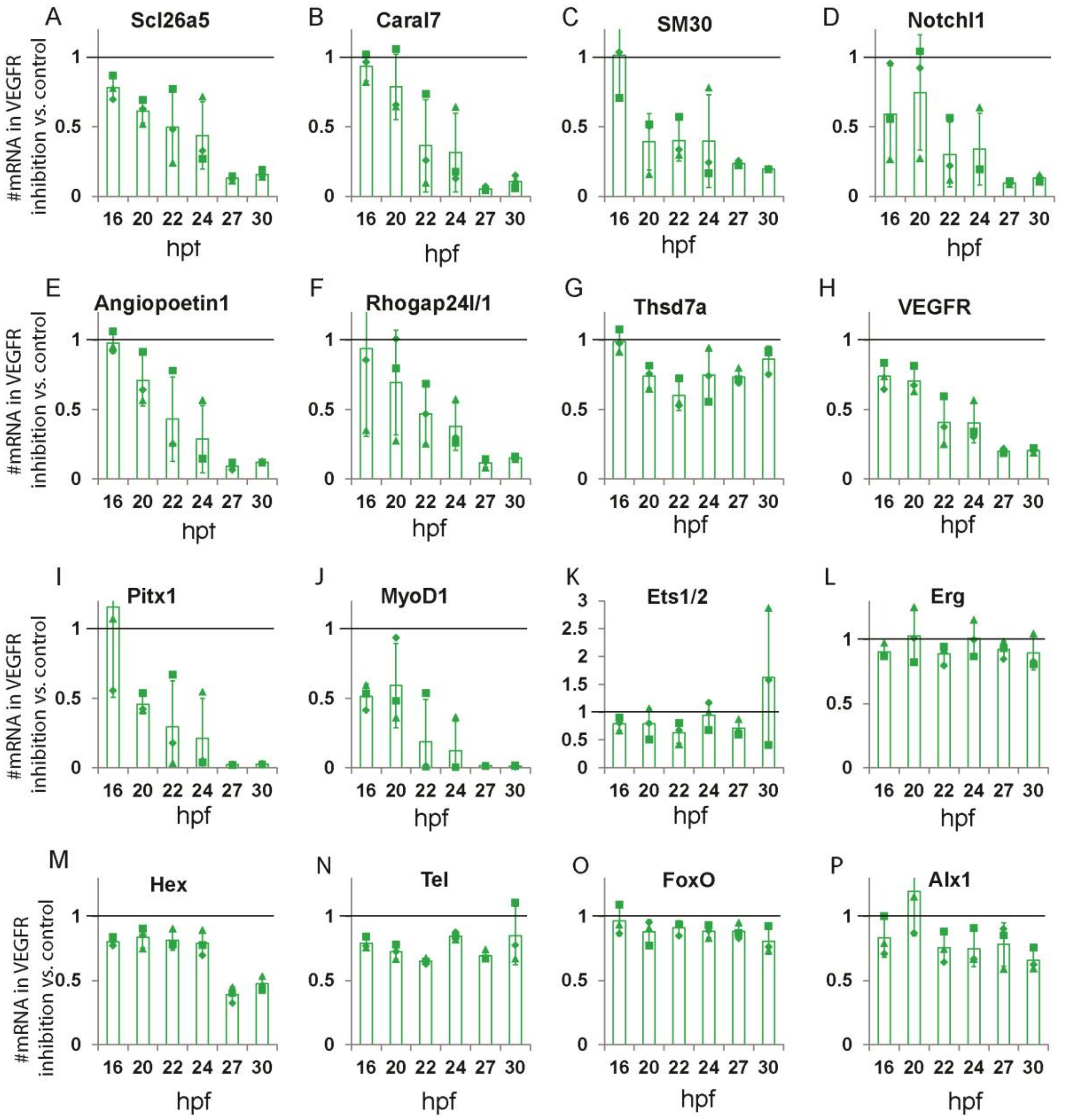


**Figure S6 Effect of VEGFR inhibition on gene expression for the genes presented in Fig. 5, 7 and S5 measured by QPCR.** Relative expression level in VEGFR inhibition vs. Control was measured by QPCR at different developmental point (n=3). Bars show averages and markers indicate individual measurement. Ratio of 1 indicates that the expression level of the gene is unaffected by VEGFR inhibition. The expression level of the genes *notchl1, rhogap24l/2, pitx1 and myoD1* is low before 22hpf therefore the results for these genes are very noisy before this time point.

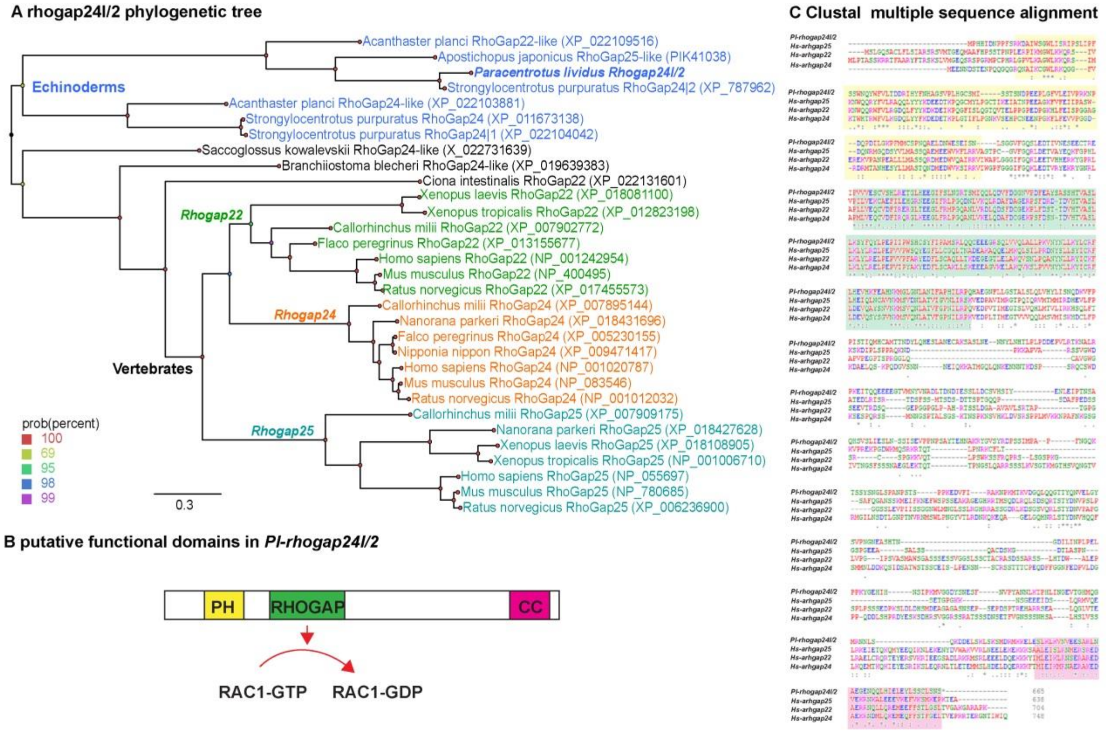


**Figure S7 *Rhogap24/2* phylogenetic tree sequence alignment and domains**. A, a phylogenetic tree of *rhogap24* gene family within deuterostomes shows that the events of gene duplications in this family occurred after the split between echinoderms and the other deuterostomes phyla (see methods for details). B, a map of *Pl*-rhogap24l/2 putative functional domains. C, sequence alignment of *Pl-rhogap24l/2*, *Hs*-*arhgap24*, *Hs*-*arhgap25* and *Hs*-*arhgap22*. Yellow background indicates the PH domain, green background the Rhogap domain highlighting the functional GAP amino acids that are conserved. Pink background indicates the CC domain.

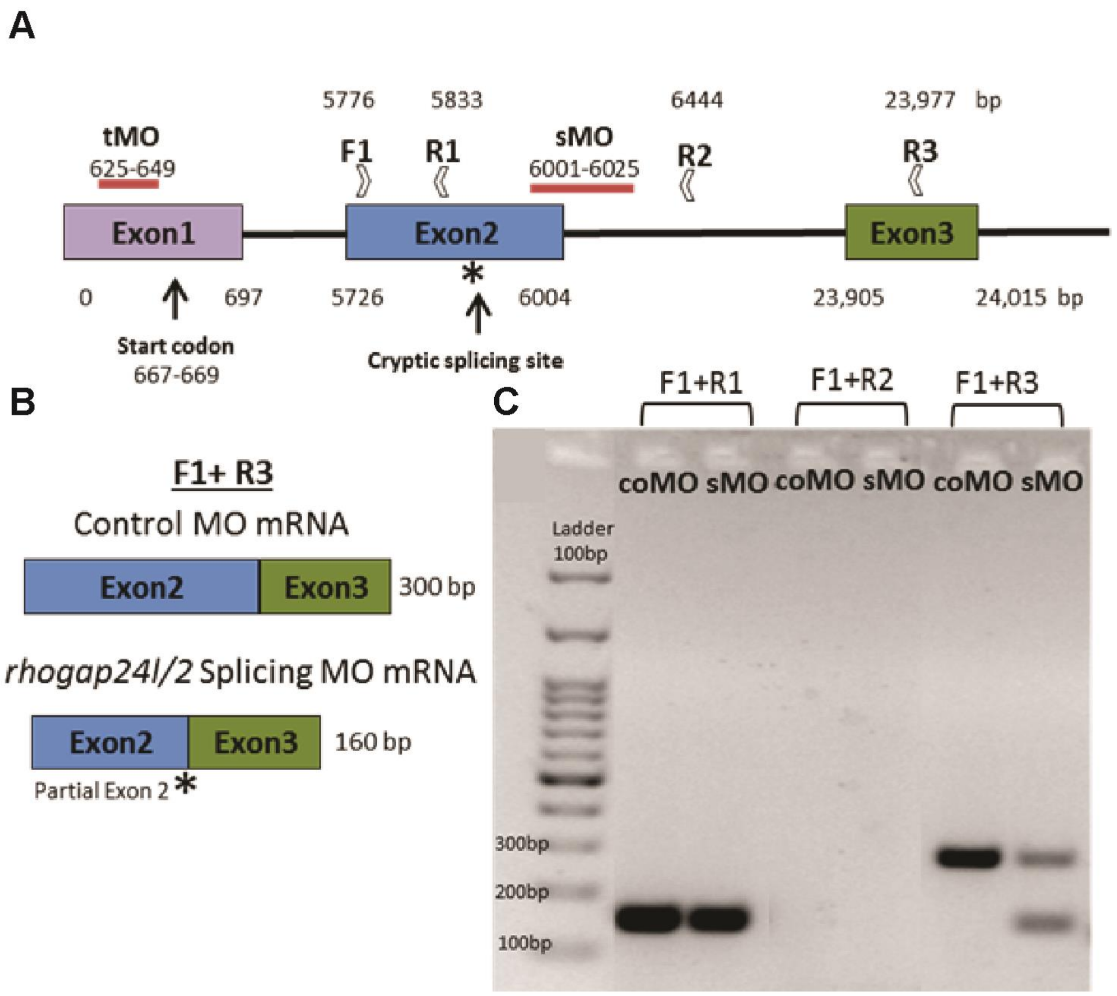


**Figure S8 Pl-*rhogap24l/2* MO design.** A, the structure of *Pl-**rhogap24l/2*** with MO target sites and primers location indicated. The coordinates correspond to the distance from the beginning of the first exon. The splicing-blocking MO (sMO) recognizes the second exon-intron boundary. The translation-blocking MO (tMO) targets a sequence positioned ~40bp upstream to the translational start codon. B, the expected PCR amplicon of the primer pair F1+R3 in control MO (coMO, normal *rhogap24l/2* transcript) and in *rhogap24l/2* splicing MO (sMO, truncated *rhogap24l/2* transcript). The truncated *rhogap24l/2* transcript (b, bottom) retains part of the second exon due to the presence of a cryptic splicing site within exon 2 (marked in asterisk in both b and a). C, PCR with different primer sets was performed at 27hpf, mid-gastrula stage, for embryos injected with 800 µM sMO and coMO. Agarose gel electrophoresis shows that the second exon is still transcribed when sMO is injected (F1+R1), splicing still occurs as the intronic primer does not produce a product in both coMO and sMO(F1+R2), but when sMO is injected *rhogap24l/2* transcript shows both normal and truncated forms (F1+R3).

**Table S1:** Number of embryos or cells scored in each experimental group in each condition. Related experiments (controls and conditions) are ordered sequentially and highlighted in similar color. For experiments #1-#7, #16-#20, numbers indicates embryos scored, for 8-15 numbers indicate cells scored.

| Exp# | Biological replicate Experimental condition | R#1 | R#2 | R#3 | R#4 | R#5 | Total number |
|---|---|---|---|---|---|---|---|
| 1 | Pl-VEGF mRNA 72hpf (Fig. 1c,d) | 43 | 28 | 43 | | | 114 |
| 2 | Hs-VEGFa(165) 72hpf (Fig. 1e,f) | 11 | 17 | 32 | 25 | | 85 |
| 3 | GFP mRNA 72hpf (Fig. 1a,b) | 35 | 23 | 24 | 32 | | 114 |
| 4 | Control MO + GFP mRNA (Fig. 1g) | 27 | 26 | 30 | | | 30 |
| 5 | VEGF MO + GFP mRNA (Fig. 1h) | 42 | 45 | 43 | | | 130 |
| 6 | VEGF MO + Pl-VEGF mRNA (Fig. 1i) | 45 | 31 | 35 | | | 111 |
| 7 | VEGF MO + Hs-VEGF mRNA (Fig. 1j) | 32 | 45 | 50 | | | 127 |
| 8 | 16hpf axitinib (Fig. 2b,c) | 21 | 21 | 25 | | | 67 |
| 9 | 16hpf DMSO (Fig. 2b,c) | 21 | 21 | 25 | | | 67 |
| 10 | 20hpf axitinib (Fig. 2b,c) | 10 | 31 | 25 | | | 66 |
| 11 | 20hpf DMSO (Fig. 2b,c) | 10 | 23 | 27 | | | 60 |
| 12 | 24hpf axitinib (Fig. 2b,c) | 10 | 32 | 29 | | | 71 |
| 13 | 24hpf DMSO (Fig. 2b,c) | 10 | 31 | 29 | | | 70 |
| 14 | 30hpf axitinib (Fig. 2b,c) | 8 | 29 | 29 | | | 66 |
| 15 | 30hpf DMSO (Fig. 2b,c) | 8 | 19 | 30 | | | 57 |
| 16 | Random MO 72hpf (Fig. 4a,f) | 33 | 37 | 45 | 36 | 58 | 209 |
| 17 | Rhogap24l/2 translation MO 72 hpf (Fig. 4b,f) | 64 | 26 | 40 | | | 130 |
| 18 | Rhogap24l/2 splicing MO 72 hpf (Fig. 4c,f) | 36 | 45 | 12 | | | 93 |
| 19 | GFP mRNA 72hpf (Fig. 4d,f) | 33 | 34 | 45 | | | 112 |
| 20 | Rhogap24l/2 mRNA 72hpf (Fig. 4e,f) | 36 | 51 | 63 | | | 150 |

**Data S1. (separate file)**

List of differentially expressed genes at VEGFR inhibition in the time course (16, 20, 24 and 30 hpf) and wash experiments (24 and 30 hpf) and both experiments combined at 24 and 30 hpf.

**Data S2. (separate file)**

List of all transcripts studied in this work with quantification at 24hpf and 30 hpf for the four replicates at 24 and 30 hpf (time course and wash combined).

**Data S3. (separate file)**

GO-enrichment within the genes differentially expressed in VEGFR inhibition at 24hpf and 30hpf.

**Data S4. (separate file)**

List of citations from which the regulatory links in vascularization were inferred to draw thick lines in Fig. 6G.

**Data S5. (separate file)**

List of primers used in this work.

7. Hu-Lowe DD, *et al.* (2008) Nonclinical antiangiogenesis and antitumor activities of axitinib (AG-013736), an oral, potent, and selective inhibitor of vascular endothelial growth factor receptor tyrosine kinases 1, 2, 3. *Clin Cancer Res* 14(22):7272-7283.
8. Bhargava P & Robinson MO (2011) Development of second-generation VEGFR tyrosine kinase inhibitors: current status. *Current oncology reports* 13(2):103-111.
9. Adomako-Ankomah A & Ettensohn CA (2013) Growth factor-mediated mesodermal cell guidance and skeletogenesis during sea urchin gastrulation. *Development* 140(20):4214-4225.
10. Grabherr MG, *et al.* (2011) Full-length transcriptome assembly from RNA-Seq data without a reference genome. *Nat Biotechnol* 29(7):644-652.
11. Haas BJ, *et al.* (2013) De novo transcript sequence reconstruction from RNA-seq using the Trinity platform for reference generation and analysis. *Nat Protoc* 8(8):1494-1512.
12. Tu Q, Cameron RA, Worley KC, Gibbs RA, & Davidson EH (2012) Gene structure in the sea urchin Strongylocentrotus purpuratus based on transcriptome analysis. *Genome Res*.
13. Li B & Dewey CN (2011) RSEM: accurate transcript quantification from RNA-Seq data with or without a reference genome. *BMC Bioinformatics* 12:323.
14. Robinson MD, McCarthy DJ, & Smyth GK (2010) edgeR: a Bioconductor package for differential expression analysis of digital gene expression data. *Bioinformatics* 26(1):139-140.
15. Robinson MD & Oshlack A (2010) A scaling normalization method for differential expression analysis of RNA-seq data. *Genome Biol* 11(3):R25.
16. Benjamini Y & Hochberg Y (1995) Controlling the false discovery rate: a practical and powerful approach to multiple testing. *J. Roy. Stat. Soc. B.* 57(1):289-300.
17. Young MD, Wakefield MJ, Smyth GK, & Oshlack A (2010) Gene ontology analysis for RNA-seq: accounting for selection bias. *Genome Biol* 11(2):R14.
18. Nakamura F (2013) FilGAP and its close relatives: a mediator of Rho-Rac antagonism that regulates cell morphology and migration. *Biochem J* 453(1):17-25.
19. Nakahara S, Tsutsumi K, Zuinen T, & Ohta Y (2015) FilGAP, a Rho-ROCK-regulated GAP for Rac, controls adherens junctions in MDCK cells. *J Cell Sci* 128(11):2047-2056.
20. Ohta Y, Hartwig JH, & Stossel TP (2006) FilGAP, a Rho- and ROCK-regulated GAP for Rac binds filamin A to control actin remodelling. *Nat Cell Biol* 8(8):803-814.
21. Su ZJ, *et al.* (2004) A vascular cell-restricted RhoGAP, p73RhoGAP, is a key regulator of angiogenesis. *Proc Natl Acad Sci U S A* 101(33):12212-12217.
22. Katoh K, Rozewicki J, & Yamada KD (2017) MAFFT online service: multiple sequence alignment, interactive sequence choice and visualization. *Brief Bioinform*.
23. Ronquist F, *et al.* (2012) MrBayes 3.2: efficient Bayesian phylogenetic inference and model choice across a large model space. *Syst Biol* 61(3):539-542.
